# Patient-derived 3D engineered human muscle model recapitulates *CLCN1* mis-splicing and myotonia in myotonic dystrophy type 1

**DOI:** 10.64898/2026.02.03.703003

**Authors:** Xiomara Fernández-Garibay, María Sabater-Arcís, Ainoa Tejedera-Villafranca, Kamel Mamchaoui, Anne Bigot, Mònica Suelves, Gisela Nogales-Gadea, Rubén Artero, Javier Ramón-Azcón, Juan M. Fernández-Costa

**Author notes:** These authors contributed equally: Xiomara Fernández- Garibay & María Sabater-Arcís.

## Abstract

Myotonic dystrophy type 1 (DM1) lacks human *in vitro* models that directly link RNA toxicity to mature skeletal muscle function, particularly myotonia. Here, we engineer contractile 3D human skeletal muscle tissues from immortalized myoblasts derived from three DM1 patients representing juvenile, adult, and late-onset subtypes. These tissues reproduce key molecular features of DM1, including nuclear RNA foci, MBNL1 sequestration, and widespread mis-splicing. Functionally, DM1 tissues exhibit impaired calcium handling, subtype-dependent weakness, rapid fatigue, and a fiber-type distribution characterized by increased slow type I fibers and pathological MyHC-I/IIx hybrids. Notably, the 3D environment enables expression and complete pathogenic mis-splicing of *CLCN1*—undetectable in matched 2D cultures—accompanied by myotonia-like delayed relaxation. Using this model, we assessed therapeutic responses of candidate small-molecule modulators. Phenylbutazone reduced RNA foci and MBNL1 sequestration but failed to rescue spliceopathy or function. In contrast, calcitriol induced coordinated transcriptomic remodeling and robustly rescued myotonia-like relaxation despite persistent *CLCN1* mis-splicing. These findings establish a functionally mature human DM1 muscle model and highlight compensatory network activation as a strategy to improve muscle function in DM1.

---

Myotonic dystrophy type 1 (DM1) is the most common inherited muscle disorder in adults and remains a chronically debilitating condition with no available cure. Clinically, it is characterized by progressive skeletal muscle wasting, weakness, and myotonia (impaired relaxation after contraction)^1^. While DM1 affects multiple organ systems, skeletal muscle impairment is one of the most functionally impactful features of the disease, leading to significant disability and reduced quality of life^2,3^.

DM1 is caused by a CTG repeat expansion in the 3′ untranslated region of the *DMPK* gene^4,5^, which leads to the production of mutant transcripts that form nuclear RNA foci and exert toxic gain-of-function effects^6^. Longer CUG expansions result in more severe symptoms in patients^7^. These expanded CUG-repeat RNAs form stable hairpin structures that sequester Muscleblind-like splicing regulator 1 (MBNL) proteins, thereby depleting them of function^8^. As a result, MBNL-regulated alternative splicing and other post-transcriptional processes are disrupted^9,10^. In parallel, CELF1 protein is abnormally stabilized, further exacerbating transcriptomic dysregulation. Therefore, at a molecular level, DM1 is characterized by a loss-of-function of MBNL and a gain-of-function of CELF1^6^.

Two-dimensional cell culture models using patient-derived fibroblasts and myoblasts were essential for understanding the molecular basis of the disease (reviewed by Matloka et al^11^). These models can recapitulate key molecular hallmarks of DM1, such as nuclear RNA foci^12–14^, mis-splicing events^15–17^, and metabolic alterations^18,19^. However, they lack the architectural and physiological complexity of native skeletal muscle, thereby limiting their ability to model DM1 functional phenotypes such as contractile weakness, fatigue, and myotonia. In contrast, mouse models ^20–27^ enable the measurement of these contractile impairments but do not accurately reflect human-specific molecular dynamics or the genetic and clinical heterogeneity of DM1.

Bridging this gap requires human-relevant systems that can integrate both molecular pathology and functional outcomes. Recently, 3D skeletal muscle models have been developed from DM1 patient-derived cells^28,29^. These 3D models support the formation of aligned myofiber filaments or sheets, which better resemble *in vivo* tissues and allow for longer culture times than 2D models. However, no functional, contractile *in vitro* model for DM1 has been developed to date. Advances in tissue engineering and microfabrication have enabled the generation of contractile myofiber bundles that allow dynamic functional measurements *in vitro*^30–37^. Applying these technologies to DM1 offers a promising platform for investigating the link between RNA toxicity and muscle dysfunction in a controlled, physiologically relevant context.

Here, we report a bioengineered 3D skeletal muscle model of DM1, generated from immortalized myoblasts derived from patients with genetic and clinical heterogeneity^38^. This is the first *in vitro* system to detect both myotonia-like impaired relaxation and *CLCN1* splicing defects in human DM1 muscle constructs. By capturing the disease’s genetic diversity and molecular hallmarks, while enabling direct measurement of functional phenotypes in a patient-specific setting, our platform offers a robust tool for mechanistic discovery and preclinical drug evaluation in DM1.

## Results

### Functional DM1 patient-derived skeletal muscle tissues exhibit weakness and fatigue reflecting disease severity across subtypes

Muscle weakness, fatigue, and myotonia are the primary functional skeletal muscle symptoms in DM1^39–42^. To model these phenotypes and reflect the disease’s clinical heterogeneity, we engineered 3D skeletal muscle tissues using immortalized myoblasts derived from three DM1 patients. These included juvenile, adult, and late-onset subtypes, each characterized by distinct CTG repeat lengths and clinical profiles, alongside three control lines from healthy donors (Supplementary Table 1^38^). As described in our previous work^43^ among the three DM1 cell lines, adult DM1-3 exhibited the highest somatic instability (>1,900 repeats) and ePAL values (>500 CTG repeats) with CTG expansions reaching up to 2,301 repeats. Juvenile DM1-1 showed comparable instability (>1,900 repeats) with peaks at 953 and 2,080 repeats, and late-onset DM1-2 presented the most moderate profile with lower instability (<700 repeats) and ePAL (<500 CTG repeats). These genetic parameters correlated directly with clinical severity based on mRS values: DM1-3 corresponded to the most severely affected patient, DM1-1 to intermediate severity, and DM1-2 to the mildest presentation, establishing DM1-3 as the cellular model with the largest CTG expansions and worst disease severity.

We encapsulated these myoblasts in a hydrogel matrix anchored between flexible posts to create tissues capable of generating measurable contractile force in response to electrical stimulation (Supplementary Fig. 1a). At day 0, all tissues appeared as uniform constructs with smooth edges (Supplementary Fig. 1b). By day 5 of differentiation, while control tissues maintained structural integrity with well-defined borders, all three DM1 lines exhibited characteristic irregular edges with bead-like protrusions (Supplementary Fig. 1c-d). After 17 days of differentiation, the tissues consisted of aligned, multinucleated myofibers expressing sarcomeric α-actinin (SAA) (Fig. 1a). At this stage, we applied electrical pulse stimulation (EPS) using a custom-built system^44^.

**Fig. 1.**
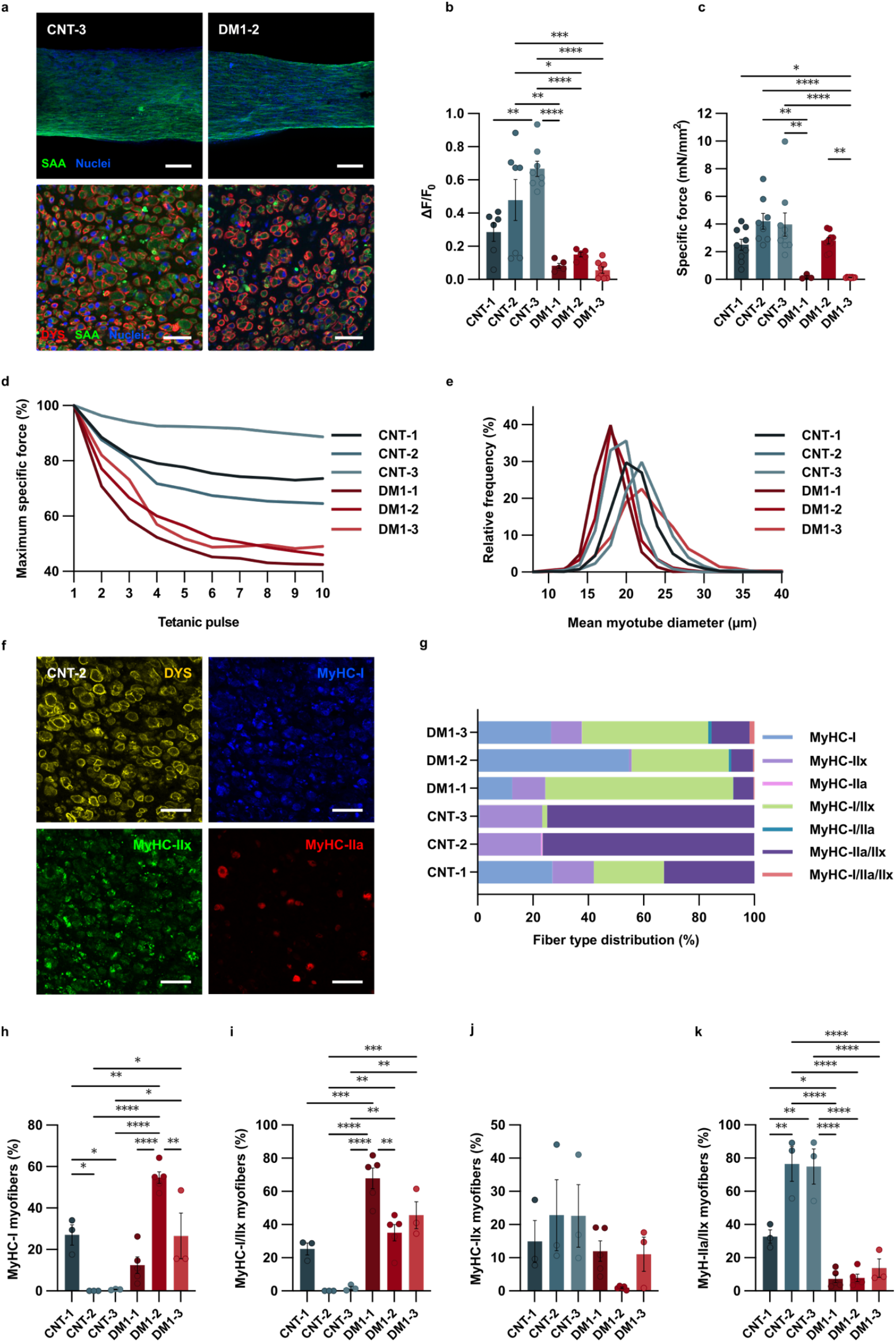
Patient-derived 3D skeletal muscle tissues recapitulate functional weakness, fatigue, and fiber-type remodeling across DM1 subtypes. **a**, Representative immunofluorescence images of control (CNT-3) and DM1 (DM1-2) skeletal muscle tissues after 17 days of differentiation. Top panels show whole-mount staining for sarcomeric α-actinin (SAA, green) and nuclei (DAPI, blue). Bottom panels show tissue cross-sections stained for dystrophin (DYS, red). Scale bars = 100 µm (top) and 40 µm (bottom). b, Maximum calcium transient amplitude (ΔF/F_0_) measured in CNT and DM1 tissues. c, Quantification of maximum specific force (mN/mm^2^). d, Normalized mean specific force (%) during repeated tetanic stimulations, revealing increased fatigability in DM1 tissues. e, Frequency distribution (%) of myotube diameters. f, Representative immunofluorescence images of tissue cross-sections stained for dystrophin (yellow) and myosin heavy chain (MyHC) isoforms: MyHC-I (blue), MyHC-IIx (green), and MyHC-IIa (red). Scale bars = 50 µm. g, Quantification of fiber-type composition showing relative percentages of MyHC-I, MyHC-IIx, MyHC-IIa, and hybrid fiber types in CNT and DM1 tissues. h-k, Individual data points and quantification of fiber-type percentages for (h) MyHC-I; (i) hybrid MyHC-I/IIx; (j) MyHC-IIx; and (k) hybrid MyHC-IIa/IIx fibers across CNT and DM1 groups. Data are shown as mean ± SEM (n ≥ 3 biological replicates per group). Statistical significance was assessed using one-way ANOVA with Tukey’s post hoc test or Student’s t-test, as appropriate. *P < 0.05; **P < 0.01; ***P < 0.001; ****P < 0.0001.

We first investigated calcium handling, a critical process for muscle contraction and a common impairment in DM1^45–48^. Calcium flux imaging during EPS revealed robust calcium transients in control tissues (ΔF/F_0_ = 0.48 ± 0.06, n = 21 pooled controls), whereas all DM1 tissues displayed significantly reduced responses (DM1-1: 0.08 ± 0.02, DM1-2: 0.15 ± 0.01, DM1-3: 0.06 ± 0.02; n = 5–8) (Fig. 1b). Overall, DM1 tissues exhibited ∼70–85% lower calcium amplitudes compared with controls, indicating a shared and severe calcium-handling defect across DM1 subtypes.

Because calcium handling and force generation can be differentially affected in DM1, we next quantified specific force across all tissues. Under high-frequency (150 Hz) tetanic stimulation, control tissues produced maximum specific forces of 3.54 ± 0.38 mN/mm^2^ (n = 26 pooled controls), whereas both juvenile (DM1-1) and adult (DM1-3) DM1 tissues generated only 0.19 ± 0.10 and 0.14 ± 0.01 mN/mm^2^, respectively (n = 3–8), corresponding to ∼95–96% reductions in strength (Fig. 1c). In contrast, late-onset DM1 tissues (DM1-2), derived from patients with fewer repeats and mild muscle involvement, produced forces comparable to controls (2.80 ± 0.23 mN/mm^2^, n = 9). This inverse relationship between CTG repeat length and contractile strength mirrors the clinical progression of DM1, linking genotype to functional muscle weakness *invitro*.

Muscle weakness alone does not explain the prominent fatigue seen in DM1. To explore this, we applied high-frequency (150 Hz) electrical stimulation for brief periods (2 s), each followed by 5 s of rest. Healthy tissues maintained their force over the stimulation regimen, declining only modestly to ∼70–85% of their initial output by the 10^th^ pulse (Fig. 1d). In contrast, all DM1 tissues—including the otherwise strong DM1-2 line—showed a pronounced drop in force immediately after the first contraction and fell to ∼40–55% of baseline by the 10^th^ pulse. This rapid loss of endurance mirrors the severe fatigability reported clinically in DM141 and recapitulated in DM1 mouse models^49^, demonstrating that our 3D system captures both muscle weakness and fatigue-like phenotypes.

To assess whether the functional deficits we observed could be attributed to structural differences, we quantified myotube diameter distributions (Fig. 1e, Supplementary Fig. 2a-b) and cross-sectional area (CSA) (Supplementary Fig. 2a-c) across all tissues. Juvenile and late-onset DM1 tissues (DM1-1 and DM1-2) formed thinner myotubes, with a mode diameter around ∼18 µm. In contrast, the adult severe subtype (DM1-3) displayed a broader diameter distribution, with the mode near ∼22 µm and diameters extending beyond 30 µm. This pattern was mirrored in the CSA analysis: DM1-1 and DM1-2 exhibited narrower distributions shifted toward smaller areas, whereas DM1-3 showed a wider range including larger fibers (Supplementary Fig. 2c). Control tissues spanned a similarly broad distribution across both metrics. Overall, differences in myotube diameter or CSA did not align with differences in specific force, indicating that atrophy-related structural changes do not account for the weakness or fatigability observed in our DM1 muscle tissues.

DM1 is often accompanied by a shift from glycolytic toward slow, oxidative type I fibers in animal models^50–52^ and patient biopsies^53–55^. To assess whether this phenotype was reproduced in our 3D tissues, we performed Myosin Heavy Chain (MyHC) immunostaining on cross-sections (Fig 1f and Supplementary Fig. 3) and quantified fiber-type distributions across all samples (Fig. 1g). All DM1 tissues exhibited markedly increased proportions of type I fibers (∼15–55%), whereas CNT-2 and CNT-3 contained almost none (∼0.5%) (Fig. 1h). DM1 tissues also displayed striking elevations in hybrid MyHC-I/IIx fibers (∼35–70% in DM1 vs ∼0.5% in CNT-2 and CNT-3) (Fig. 1i). These hybrid fibers are rarely detected in healthy human or mouse muscle and are considered pathological indicators of fiber-type disorganization^56–58^. Notably, CNT-1, a control line that naturally forms a higher proportion of oxidative fibers, showed MyHC-I and MyHC-I/IIx levels comparable to some DM1 tissues. In contrast, control tissues were dominated by fast glycolytic MyHC-IIx fibers (∼15–23%) (Fig. 1j) and by physiologically common hybrid IIa/IIx fibers (∼32–76% in controls vs ∼7–13% in DM1) (Fig. 1k), consistent with healthy human muscle^58–60^. Collectively, these analyses reveal a pronounced shift toward oxidative and pathological hybrid fiber types in DM1 tissues.

Together, these results show that our patient-derived muscle constructs accurately recapitulate core hallmarks of DM1—including impaired calcium dynamics, contractile weakness, rapid fatigue, and a shift toward oxidative fiber identity—demonstrating that fiber-type dysregulation is a robust and quantifiable feature of the disease in engineered human muscle.

### Patient-derived muscle tissues exhibit hallmark molecular defects

The functional deficits observed in our bioengineered DM1 tissues, from impaired contractility to altered fiber composition, reflect underlying molecular abnormalities characteristic of DM1. To determine whether our patient-derived 3D skeletal muscle tissues recapitulate these molecular hallmarks, we first examined the presence of intranuclear RNA foci, a defining feature of the disease caused by expanded CUG repeats in the 3’ UTR of *DMPK* transcripts^6^. Using fluorescence in situ hybridization (FISH), we detected abundant RNA foci exclusively in cryosections from DM1 tissues, with none observed in controls (Fig. 2a, Supplementary Fig. 4). The number of foci per nucleus varied significantly across subtypes, consistent with CTG repeat lengths: DM1-1 (juvenile) and DM1-3 (adult) tissues exhibited the highest foci burden (∼1.5-1.6 foci per nucleus), whereas DM1-2 (late-onset) showed fewer foci (∼1.1 foci per nucleus) (Fig. 2b). Notably, DM1-2 tissues, those with fewer RNA foci, also showed the strongest contractile performance, suggesting a potential relationship between RNA toxicity and functional impairment.

**Fig. 2.**
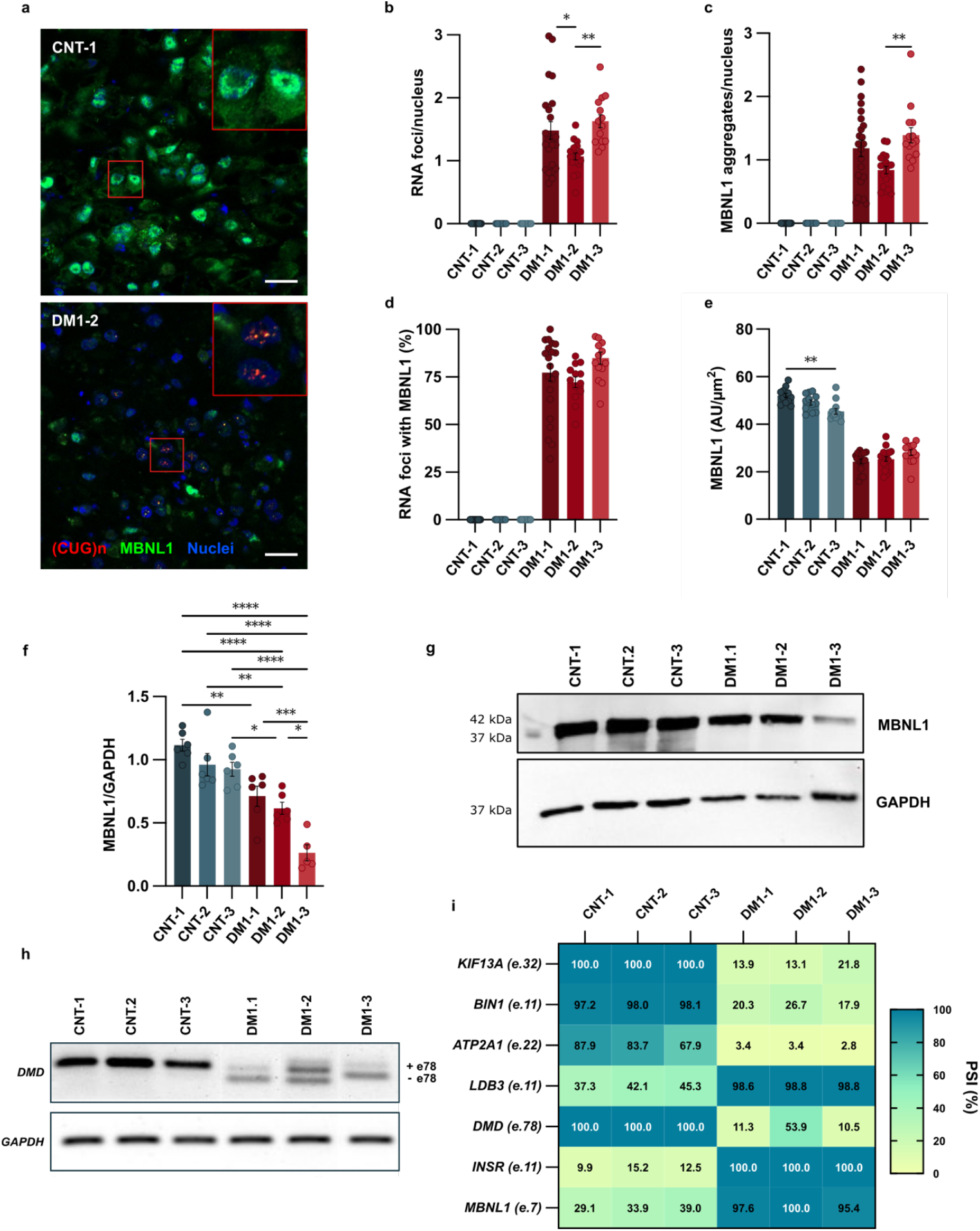
Patient-derived 3D skeletal muscle tissues exhibit key molecular hallmarks of DM1. **a**, Representative confocal images of ribonuclear RNA foci and MBNL1 localization in transversal cross-sections of control (CNT-1) and DM1-2 tissues after 17 days of differentiation. (CUG)n RNA foci were detected by fluorescence *in situ* hybridization using a Cy3-labelled probe (red), MBNL1 was immunostained (green), and nuclei were counterstained with DAPI (blue). Scale bars = 20 µm. b-e, Quantification of (b) RNA foci per nucleus, (c) MBNL1 aggregates per nucleus, (d) percentage of RNA foci colocalizing with MBNL1, and (e) corrected total cell fluorescence of MBNL1 (AU/µm^2^). A total of 12-18 images per condition were quantified. In panels (b-e), the global CNT versus DM1 comparison was highly significant (P < 0.0001, one-way ANOVA) and is not shown on the graphs for clarity. f-g, Western blot (f) quantification and (g) representative blot images of MBNL1 expression across control and DM1 tissues. *GAPDH* was used as an endogenous control. h-i, Analysis of alternative splicing by semiquantitative RT–PCR. (h) Representative gel images for *DMD* (exon 78) and *GAPDH*. (i) Heatmap showing percent spliced-in (PSI, %) values for *KIF13A* (exon 32), *BIN1* (exon 11), *ATP2A1* (exon 22), *LDB3* (exon 11), *DMD* (exon 78), *INSR* (exon 11), and *MBNL1* (exon 7) across CNT and DM1 3D muscle tissues. The bar graphs show mean ± SEM. \**P* < 0.05; \*\**P* < 0.01; \*\*\**P* < 0.001 and \*\*\*\**P* < 0.0001 by a one-way ANOVA test and Tukey’s HSD post hoc test.

Next, we assessed whether these RNA foci sequestered MBNL1, a key alternative splicing regulator. Confocal microscopy of immunostained sections revealed nuclear MBNL1 aggregates that co-localized with RNA foci in all DM1 lines, but not in controls (Fig. 2a). Quantitative analysis demonstrated MBNL1 aggregates in all DM1 lines (DM1-1: ∼1.2, DM1-2: ∼0.9, DM1-3: ∼1.4 aggregates per nucleus) compared to none in controls (Fig. 2c). The percentage of RNA foci containing MBNL1 was likewise elevated in DM1 tissues (DM1-1: ∼77%, DM1-2: ∼72%, DM1-3: ∼85%) (Fig. 2d). Furthermore, MBNL1 fluorescence intensity was significantly reduced in both the cytoplasm and nuclei of DM1 tissues (Fig. 2e), causing reduced functional MBNL1 availability.

Consistent with previous reports demonstrating MBNL1 protein reduction in DM1^43,61^, Western blot analysis revealed significantly lower MBNL1 protein levels in all DM1 3D muscle models compared to controls (Fig. 2f-g). Specifically, DM1-1 showed a 28.5% reduction (1.40-fold), DM1-2 demonstrated a 38.4% reduction (1.62-fold), and DM1-3 displayed the most severe reduction, around 77.2% (4.38-fold) compared to the control average. These results suggest that reduced MBNL1 protein levels correlate with the size of repeat expansion in these patient-derived 3D muscle models. Together, the repeat-dependent reduction in MBNL1 protein and the corresponding RNA foci burden establish a clear molecular pattern consistent with the clinical severity of the donor subtypes.

MBNL1 sequestration and depletion are known to drive widespread mis-splicing of transcripts critical to muscle function^8^. We found that bioengineered DM1 tissues exhibited mis-splicing in several genes, including those involved in intracellular transport (*KIF13A* exon 32), membrane remodeling (*BIN1* exon 11), calcium handling (*ATP2A1* exon 22), sarcomere structure (*LDB3* exon 11), dystrophin (DMD exon 78), insulin regulation (*INSR* exon11) and *MBNL1* itself (exon 7). Splicing analysis revealed significant alterations in percent spliced inclusion (PSI) across all DM1 lines compared with controls, with gene-specific variability (Figure 2h-i, Supplementary Fig. 5). Notably, *DMD* displayed an intermediate PSI alteration (∼53.9%) in DM1-2 relative to the more severe changes observed in DM1-1 and DM1-3 (PSI -11.3% and ∼10.5% respectively), consistent with the late-onset clinical form and reduced disease severity in DM1-2 patient. Overall, these widespread splicing defects confirm that the molecular and functional abnormalities caused by toxic RNA can be replicated in our *in vitro* model.

### Transcriptomic analysis reveals widespread dysregulation of muscle-related gene expression in DM1 tissues

While targeted analyses confirmed hallmark molecular defects, including RNA foci formation, MBNL1 sequestration, and mis-splicing of key transcripts, these alterations represent only a subset of the molecular disruptions underlying DM1. To obtain a more comprehensive view of disease-associated changes, we performed bulk RNA sequencing of control and DM1-derived skeletal muscle tissues.

Multidimensional scaling (MDS) analysis revealed a clear separation between DM1 and control samples. The late-onset and adult subtypes (DM1-2 and DM1-3) clustered closely together. In contrast, the juvenile subtype (DM1-1) exhibited greater transcriptomic divergence (Fig. 3a), highlighting the molecular heterogeneity of DM1 and the importance of capturing this diversity in *in vitro* models.

**Fig. 3.**
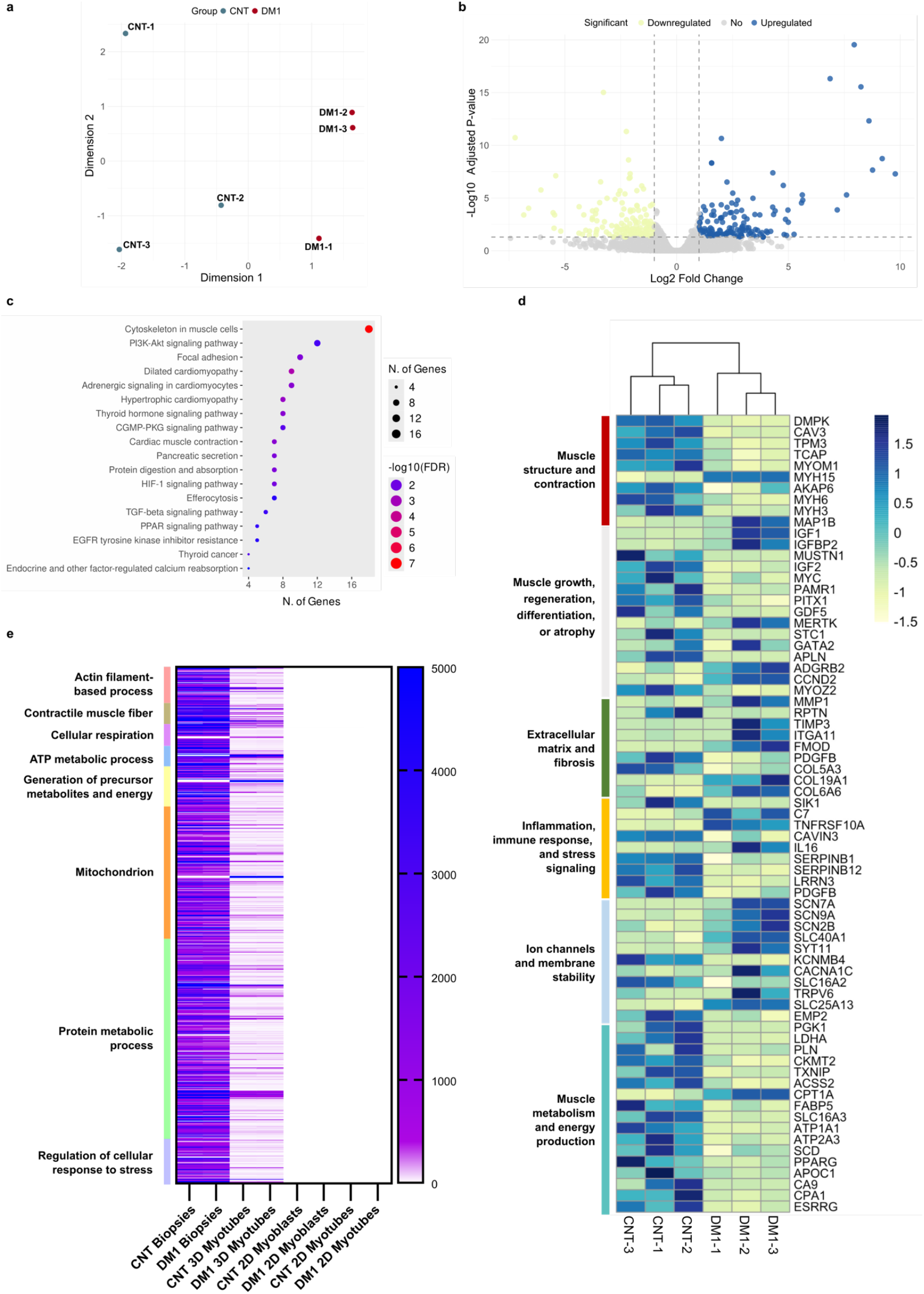
Transcriptomic dysregulation of muscle-related gene programs in DM1 3D tissues. **a**, Multidimensional scaling (MDS) plot of RNA-seq profiles from control (CNT) and DM1 engineered muscle tissues differentiated for 17 days. b, Volcano plot of differentially expressed genes (DEGs) between CNT and DM1. c, Heatmap of selected statistically significant DEGs identified by RNA-seq, clustered according to functional gene categories. d, Gene ontology enrichment analysis of differentially expressed genes using KEGG pathway annotation. e, Functional annotation clustering of genes expressed in human muscle biopsies and 3D muscle tissues but not detected in matched 2D cultures reveals enrichment of categories related to actin filament–based processes, contractile muscle structure, energy metabolism, cellular respiration, mitochondrial function, protein metabolism, and regulation of cellular stress responses. RNA-seq analysis was performed using n = 3 biological replicates per condition; each biological replicate comprised 6–8 technical replicates. Differential expression thresholds were set at |log2 FC| ≥ 1 with adjusted P < 0.05.

Differential expression analysis identified 366 genes that were significantly dysregulated in DM1 tissues compared to controls (Fig. 3b). Transcripts downregulated in DM1 tissues included genes involved in sarcomere structure and contractility *(CAV3, TPM3, MYOM1, MYH6, MYH3)*, myogenic growth and differentiation *(MUSTN1, IGF2, MYC)*, and metabolic regulation, including energy production *(ESRRG, ATP1A1, ATP2A3)*. In contrast, genes associated with inflammation *(IL16, C7, TNFRSF10A)*, and fibrosis *(FMOD, COL19A1, COL6A6, MMP1)* were upregulated (Fig. 3c).

Gene Ontology (GO) enrichment analysis supported these findings, revealing significant overrepresentation of pathways related to muscle contraction mechanics, calcium ion transport and homeostasis, and extracellular matrix remodeling (Fig. 3d, Supplementary Fig. 6). The most enriched biological processes encompassed myofibrillar differentiation, cell developmental signaling, and inflammatory response modulation, while cellular components highlighted aberrant ECM composition, plasma membrane organization, and synaptic architecture. Molecular functions centered on protein-containing complex binding, calcium-dependent signaling, and monatomic ion transmembrane transport. Together, these transcriptomic alterations demonstrate widespread impairments in structural integrity, metabolism, and ion homeostasis—hallmark molecular defects that collectively underpin DM1 muscle pathophysiology in 3D tissue models.

To evaluate the degree of functional maturation achieved in our 3D muscle tissues, we compared their transcriptomic profiles with 2D cultures and human muscle biopsies using published reference datasets^62^. We identified a subset of genes that were silent in 2D cultures but expressed in both our 3D constructs and native muscle tissue, indicating acquisition of muscle-specific functional programs. GO analysis revealed strong enrichment for pathways associated with sarcomere assembly, contractile fiber organization, mitochondrial structure and metabolism, and cellular component biogenesis, features characteristic of mature skeletal muscle (Supplementary Fig. 7a-b). Consistent with this, 3D tissues exhibited upregulation of key contractile and cytoskeletal components as well as genes involved in oxidative phosphorylation and mitochondrial function (Fig. 3e, Supplementary Fig. 7c). Together, these findings demonstrate that 3D culture promotes transcriptional programs consistent with advanced muscle maturation, providing greater physiological relevance than conventional 2D systems.

To determine how these molecular perturbations contribute to specific functional hallmarks of DM1, we next assessed whether our model exhibits myotonia-like contractile behavior and ion channel dysregulation.

### First *in vitro* recapitulation of myotonia-like relaxation defects and *CLCN1* splicing abnormalities in a DM1 model

Myotonia, characterized by impaired muscle relaxation following contraction, is a hallmark clinical manifestation of DM1^63^. To determine whether our bioengineered DM1 muscle tissues reproduced this phenotype, we analyzed relaxation kinetics from the time-normalized specific force traces acquired during 15 brief tetanic stimulation periods separated by resting intervals. While control tissues consistently returned to baseline force after each EPS period, DM1-derived tissues exhibited prolonged contraction after stimulation had ceased (Fig. 4a,b), indicative of myotonia-like behavior. We quantified this relaxation defect by calculating the integrated area under the curve (AUC) during the post-stimulation resting phases. Control tissues showed low AUC values (CNT-1: 2.90 ± 0.45; CNT-2: 2.05 ± 0.20; CNT-3: 1.30 ± 0.26), whereas all DM1 tissues displayed significantly elevated myotonia-like AUC values (Fig. 4c). The strongest defect was observed in DM1-1 (16.30 ± 2.78), DM1-2 exhibited intermediate levels (8.08 ± 0.78), and DM1-3 showed milder but still increased myotonia (3.02 ± 0.35). These differences reflect the intrinsic heterogeneity of the patient-derived lines captured by the model, without following a strict relationship with CTG repeat length or clinical severity. Notably, this represents the first quantitative assessment of myotonia-like behavior in a patient-derived 3D muscle tissue model, providing functional validation of DM1 pathophysiology in an engineered human system.

**Fig. 4.**
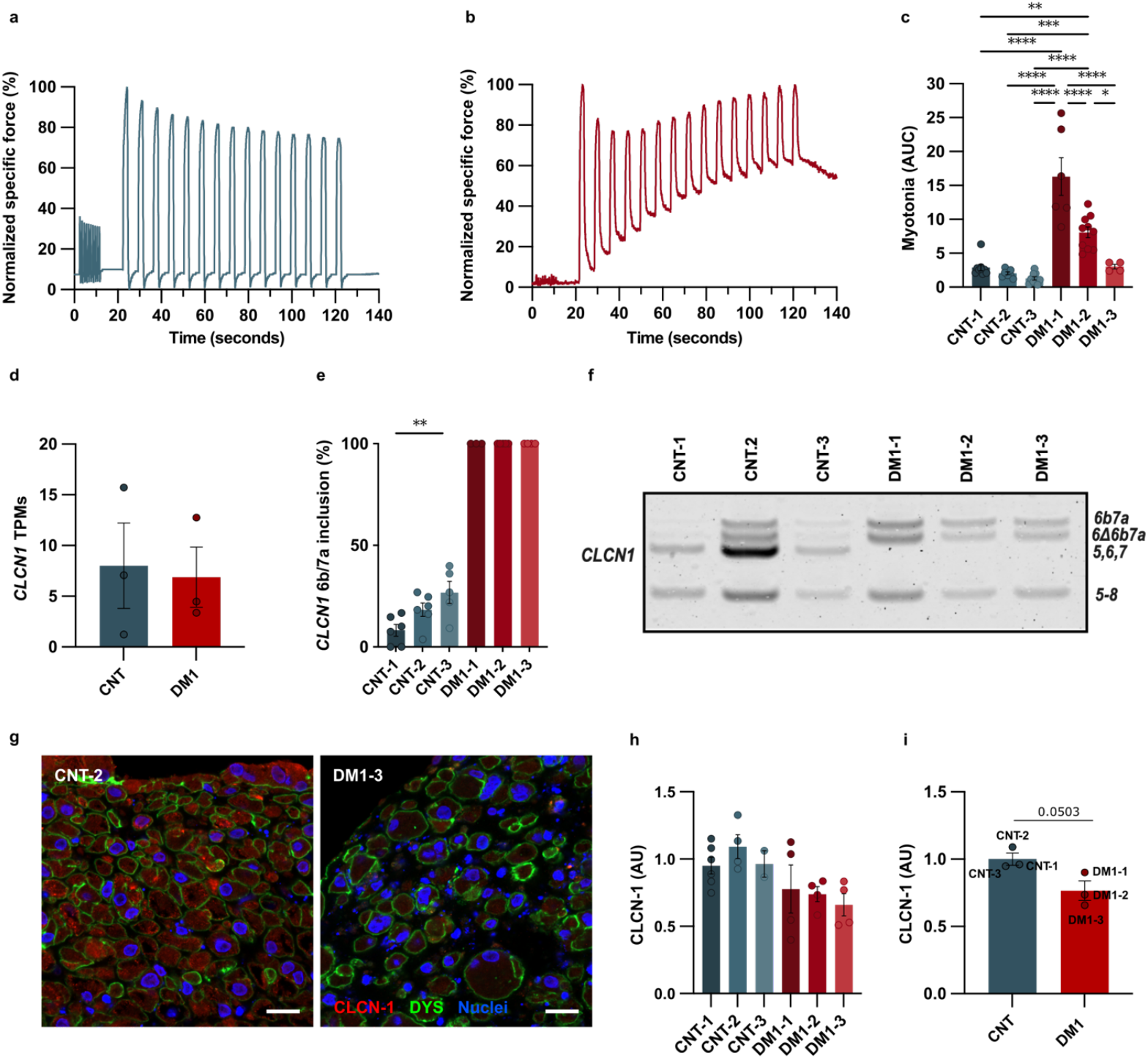
Detection of myotonia and *CLCN1* mis-splicing in engineered human DM1 muscle tissues. **a-b**, Representative traces of normalized specific force (%) relative to the maximum force in (a) CNT and (b) DM1 engineered muscle tissues. c, Quantification of myotonia as the area under the curve (AUC) of normalized specific force (%) traces during relaxation phases. d, *CLCN1* transcript abundance measured as transcripts per million (TPM) from RNA-seq analysis of CNT and DM1 3D muscle tissues; each point represents one cell line. e, Percentage of exon inclusion for *CLCN1* exons 6b/7a assessed by semi-quantitative RT–PCR in CNT and DM1 tissues; *GAPDH* was used as an internal control. The global CNT versus DM1 comparison was highly significant (P < 0.0001, one-way ANOVA) and is not shown on the graph for clarity. f, Representative agarose gel used for quantification shown in (e). g, Representative confocal images of CLCN-1 protein (red), dystrophin (DYS, green), and nuclei (DAPI, blue) in CNT and DM1. Scale bars = 20 µm. h-i, Quantification of CLCN-1 protein expression from confocal images shown in (g). (h) Corrected total cell fluorescence (AU) of CLCN-1 within myotubes and (i) mean values (AU) aggregated per CNT and DM1 cell line. Data are presented as mean ± SEM. A total of 5–6 images were quantified per cell line for immunofluorescence analyses. Statistical significance was assessed using one-way ANOVA with Tukey’s post hoc test (c,e, h) and unpaired two-tailed Student’s t-test (d,i). **P < 0.01, ****P < 0.0001.

Myotonia in DM1 has been causally linked to aberrant alternative splicing of the skeletal muscle chloride channel gene, *CLCN1*, resulting in the aberrant inclusion of exons 6b and 7a. This splicing defect introduces premature stop codons, leading to truncated protein products that are rapidly degraded by nonsense-contraction after stimulation had ceased (Fig. 4a,b), indicative of myotonia-like behavior. We quantified this relaxation defect by calculating the integrated area under the curve (AUC) during the post-stimulation resting phases. Control tissues showed low AUC values (CNT-1: 2.90 ± 0.45; CNT-2: 2.05 ± 0.20; CNT-3: 1.30 ± 0.26),whereas all DM1 tissues displayed significantly elevated myotonia-like AUC values (Fig. 4c). The strongest defect was observed in DM1-1 (16.30 ± 2.78), DM1-2 exhibited intermediate levels (8.08 ± 0.78), and DM1-3 showed milder but still increased myotonia (3.02 ± 0.35). These differences reflect the intrinsic heterogeneity of the patient-derived lines captured by the model, without following a strict relationship with CTG repeat length or clinical severity. Notably, this represents the first quantitative assessment of myotonia-like behavior in a patient-derived 3D muscle tissue model, providing functional validation of DM1 pathophysiology in an engineered human system.

Myotonia in DM1 has been causally linked to aberrant alternative splicing of the skeletal muscle chloride channel gene, *CLCN1*, resulting in the aberrant inclusion of exons 6b and 7a. This splicing defect introduces premature stop codons, leading to truncated protein products that are rapidly degraded by nonsense-mediated decay (NMD), ultimately resulting in loss of functional chloride channel^64^. Given the robust myotonia-like phenotype observed, we examined our RNA-seq data and found that *CLCN1* transcripts were expressed in 3D constructs (Fig. 4d), indicating that the 3D environment promotes the acquisition of a mature splicing program. To confirm this, we performed RT-PCR on the 3D tissues and quantified exon 6b/7a inclusion. Control tissues showed low and variable levels of pathogenic exon inclusion (CNT-1: 9.8 ± 3.2%; CNT-2: 17.9 ± 3.1%; CNT-3: 26.6 ± 3.8%), whereas all DM1 tissues exhibited complete inclusion of exons 6b/7 (Fig. 4e-f). At the protein level, immunofluorescence staining demonstrated reduced CLCN-1 signal in 3D DM1 tissues compared with robust CLCN-1 expression in controls (Fig. 4g-i, Supplementary Fig. 8), corroborating the splicing defect’s impact on channel abundance.

To verify that this mis-splicing and expression pattern is unique to our 3D model, we differentiated the same DM1 and control myoblasts in 2D under identical conditions. After 17 days, 2D myotubes showed no detectable *CLCN1* transcripts, despite robust *GAPDH* amplification, confirming that this aberrant *CLCN1* splice pattern occurs exclusively in the 3D model (Supplementary Fig. 9). This constitutes the first demonstration of *CLCN1* mis-splicing and corresponding myotonia-like physiology in a patient-derived *in vitro* DM1 muscle model, underscoring the physiological maturity of our 3D model and its capacity to reproduce both functional and molecular hallmarks of the disease.

### Treatment with known MBNL1 upregulators reduces RNA foci but fails to rescue spliceopathy in patient-derived tissues

Given the robust recapitulation of DM1 molecular and functional hallmarks in our 3D model and the high degree of physiological maturity achieved, we next investigated whether these pathological features could be therapeutically targeted to validate its utility as a preclinical drug-testing platform. Because MBNL1 functional depletion drives DM1-associated splicing dysregulation (Fig. 2), we evaluated small molecules previously reported to increase MBNL1 levels.

We initially treated DM1-3 tissues, the most severely affected line, with four compounds: phenylbutazone (324 µM) ^65,66^, calcitriol (1 µM) ^67^, vorinostat (10 µM) ^68^, and chloroquine (10 µM) ^69^, to assess their impact on contractile function. Vorinostat and chloroquine significantly impaired muscle contractility, showing a reduced calcium transient response compared to untreated (DMSO 0.01%) DM1-3 tissues (Supplementary Fig. 10), and were therefore excluded from further analysis. We subsequently focused on phenylbutazone (PBZ) and calcitriol (CAL), both of which preserved contractile function, and extended these treatments to all DM1 tissues.

At the molecular level, FISH and MBNL1 immunostaining revealed that both compounds significantly reduced RNA foci in the most affected lines (Fig 5a). In DM1-1 tissues, PBZ lowered foci from ∼1.45 to ∼0.88 foci/nucleus and CAL to ∼0.76. Similarly, in DM1-3 tissues, PBZ decreased foci from ∼1.56 to ∼0.99 foci/nucleus and CAL reduced them to ∼1.2 (Fig. 5a-b, Supplementary Fig. 11). In contrast, DM1-2 tissues showed no significant changes, consistent with their lower baseline foci burden.

**Fig. 5.**
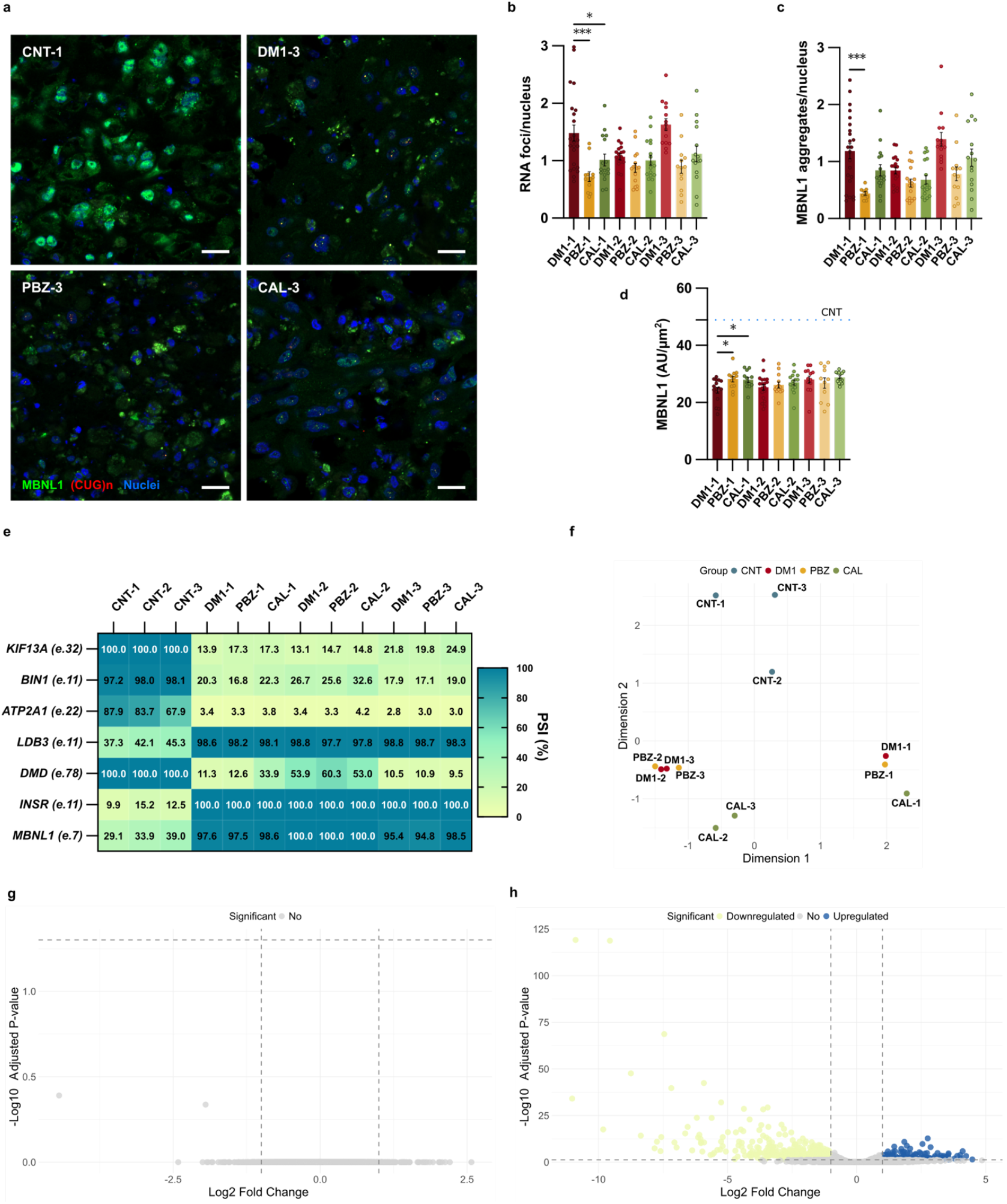
Differential effects of phenylbutazone and calcitriol on MBNL1 sequestration, splicing, and transcriptomic profiles in DM1 muscle tissues. **a**, Representative confocal images of transverse sections showing MBNL1 immunostaining (green) and (CUG)n RNA foci detected by fluorescence in situ hybridization (Cy3-labelled probe, red) in CNT-1, DM1-3, and DM1-3 tissues treated with phenylbutazone (PBZ) or calcitriol (CAL). Nuclei were counterstained with DAPI (blue). Scale bars = 20 µm.b-d, Quantification of (b) RNA foci per nucleus, (c) MBNL1 aggregates per nucleus, and (d) corrected total cell fluorescence of MBNL1 (AU/um^2^). A total of 12–18 images were quantified per condition. e, Heatmap showing percentage spliced-in (PSI, %) values for selected DM1-relevant alternative splicing events, assessed by semi-quantitative RT–PCR in untreated DM1 tissues and DM1 tissues treated with PBZ or CAL. f, Principal component analysis (PCA) of RNA-seq data from control (CNT), untreated DM1, PBZ-treated DM1, and CAL-treated DM1 3D muscle tissues. g,h, Volcano plots of differentially expressed genes comparing (g) untreated DM1 versus PBZ-treated DM1 tissues, and (h) untreated DM1 versus CAL-treated DM1 tissues. Data are presented as mean ± SEM. Statistical significance was assessed using one-way ANOVA followed by Tukey’s HSD or Dunnett’s post hoc test, comparing each treatment condition to the corresponding untreated DM1 line. *P < 0.05; **P < 0.01; ***P < 0.001; ****P < 0.0001.

PBZ also significantly decreased MBNL1 colocalization with RNA foci, reducing MBNL1 aggregates per nucleus from ∼1.15 to ∼0.63 in DM1-1 and from ∼1.43 to ∼0.77 in DM1-3, indicating reduced protein sequestration. However, CAL had no detectable effect on MBNL1 localization patterns in any DM1 line (Fig. 5c). Total MBNL1 fluorescence intensity levels remained essentially unchanged across treatments, with only modest increases in DM1-1 tissues following either PBZ or CAL treatments (Fig. 5d). Despite these molecular improvements, neither treatment rescued the spliceopathy. Splicing defects persisted across multiple MBNL1-regulated transcripts (Fig. 5e, Supplementary Fig. 5), indicating that partial reductions in RNA foci and MBNL1 sequestration were insufficient to restore normal splicing patterns in our 3D DM1 muscle model.

To further assess the therapeutic impact at the transcriptomic level, we performed RNA-seq analysis of all DM1 tissues treated with PBZ and CAL and compared them with untreated DM1 tissues. RNA-seq analysis revealed no substantial rescue of global gene expression patterns following treatments. In multidimensional scaling (MDS) plots, PBZ- and CAL-treated samples remained clearly separated from controls, with no shift toward a healthy control-like expression profile (Fig. 5f). Differential gene expression analysis further confirmed that PBZ had minimal transcriptomic impact, with no genes significantly altered compared to untreated DM1 tissues (Fig. 5g).

In contrast, CAL induced widespread transcriptomic changes, with 685 altered genes relative to untreated DM1 tissues (Fig. 5h). However, these changes did not shift CAL-treated tissues toward a control-like profile and instead reflected broad modulation of unrelated cellular pathways,

Together, these findings suggest that while PBZ partially reduces MBNL1 sequestration with minimal disruption to the transcriptome, neither compound reverses DM1-associated molecular pathology in our 3D muscle model. The extensive transcriptomic shifts induced by CAL, despite having no effect on MBNL1 localization or splicing, suggest activation of alternative cellular pathways that may confound therapeutic interpretation and require further investigation.

### Calcitriol engages compensatory pathways that improve DM1 muscle function

Since calcitriol treatment did not affect MBNL1 expression despite reported efficacy in DM1 models^67^, we first verified whether CAL was biologically active in our 3D model. Transcriptomic analysis revealed robust activation of 10vitamin D receptor (VDR)–dependent transcription, including strong induction of canonical vitamin D target genes such as *CYP24A1*, together with enrichment of vitamin D–responsive biological processes (Supplementary Fig. 12a–c). These data confirm effective VDR engagement and demonstrate that CAL is biologically active in 3D DM1 muscle tissues.

Having confirmed robust VDR activation, we next examined the broader transcriptomic profile to understand the effects of CAL on DM1 pathophysiology. Functional gene clustering analysis showed coordinated upregulation of genes involved in muscle structure and contraction, muscle growth and regeneration, and ion channels critical for muscle stability (Fig. 6a). CAL treatment also induced downregulation of pathways associated with extracellular matrix remodeling and inflammatory or stress responses, which are typically elevated in DM1, while broadly upregulating programs related to muscle metabolism and calcium homeostasis. These transcriptomic changes are consistent with improved bioenergetic capacity and normalization of dysregulated calcium signaling (Supplementary Fig. 13).

**Fig. 6.**
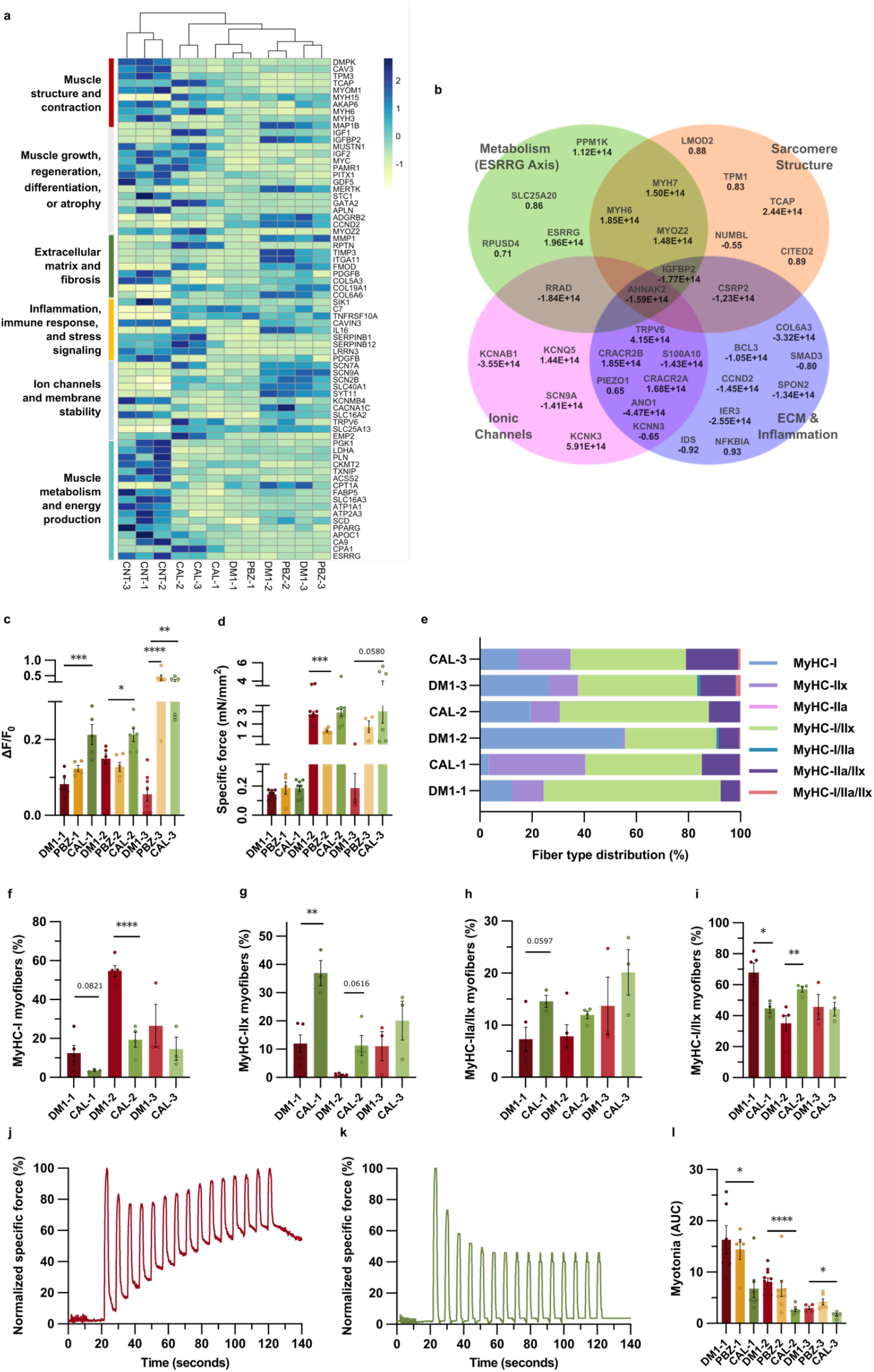
Calcitriol alleviates myotonia in patient-derived DM1 tissues independent of *CLCN1* splicing defects and induces functional and structural remodeling. **a**, Heatmap of significantly differentially expressed genes (DEGs) from RNA-seq comparing CNT, untreated DM1, and CAL-treated DM1 3D muscle tissues, clustered by functional categories. b, Venn diagram showing DEGs (fold change) in CAL-treated versus untreated DM1 tissues across four functional modules: metabolic pathways, sarcomeric structure, ion channels, and extracellular matrix (ECM)/inflammatory signaling. Intersections indicate genes modulated across multiple functional categories. c, Maximum calcium transient amplitude (ΔF/F_0_) measured in the untreated DM1 lines and treated with PBZ or CAL. d, Quantification of maximum specific force (mN/mm^2^). e, Quantification of fiber-type composition showing relative percentages of MyHC-I, MyHC-IIx, MyHC-IIa, and hybrid fiber types in untreated DM1, PBZ- and CAL-treated tissues. f-i, Individual data points and quantification of selected fiber-type populations: (f) MyHC-I, (g) MyHC-IIx, (h) MyHC-IIa/IIx, and (i) MyHC-I/IIx) in untreated DM1 and CAL-treated tissues. j-k, Representative traces of normalized specific force (%) relative to the maximum force in (j) DM1 and (k) CAL-treated tissues. l, Quantification of myotonia measured as the area under the curve (AUC) of the normalized specific force (%) during relaxation phases in untreated DM1, PBZ-, and CAL-treated tissues. Statistical analyses were performed using one-way ANOVA with Dunnett’s post hoc test for comparisons of treated versus untreated DM1 conditions. Bar graphs show mean ± SEM (n ≥ 3 biological replicates per group). Significance levels: *P < 0.05; **P < 0.01; ***P < 0.001 and****P < 0.0001.

Differential expression analysis revealed that calcitriol engages four interconnected compensatory molecular programs: restoration of mitochondrial metabolic control, reconstruction of sarcomeric architecture, suppression of the pro-fibrotic extracellular matrix (ECM), and ionic compensation of membrane excitability (Fig. 6b), the latter examined in a subsequent subsection.

In line with this framework, CAL induced upregulation of metabolic gene programs that are suppressed in DM1, including increased expression of *ESRRG*, a key regulator of mitochondrial oxidative metabolism. This response was accompanied by coordinated upregulation of mitochondrial genes involved in energy production and substrate transport, consistent with enhanced metabolic capacity in treated tissues. In parallel, CAL enhanced expression of structural genes associated with sarcomeric organization and contractile stability, including recovery of Z-disc– and thin-filament–associated components that are downregulated in untreated DM1 tissues, indicating partial reinforcement of the contractile apparatus and a shift toward a more mature muscle phenotype. Calcitriol also modulated extracellular matrix and inflammatory gene programs dysregulated in DM1 muscle, including downregulation of *COL6A3* and other profibrotic mediators, alongside reduced expression of inflammatory regulators and the DM1-associated marker *IGFBP2*. Increased expression of the anti-inflammatory factor *NFKBIA* and suppression of cell-cycle and lysosomal genes altered in DM1 were consistent with partial normalization of tissue homeostasis (Fig. 6b).

These transcriptomic and molecular changes coincided with functional improvements. Calcitriol enhanced calcium handling across all DM1 subtypes (Fig. 6c). In DM1-1 tissues, CAL increased ΔF/F_0_ from ∼0.10 to ∼0.21. DM1-2 tissues showed a similar improvement (∼0.16 → ∼0.25). The strongest effect was observed in DM1-3 tissues, where CAL increased ΔF/F_0_ from ∼0.09 to ∼0.40, representing the largest calcium-handling enhancement among all subtypes.

Despite these improvements, maximum specific force remained unchanged, with no statistically significant effects observed in any DM1 line (Fig. 6d). In DM1-1 tissues, CAL produced no meaningful recovery in force (∼0.15 to ∼0.20 mN/mm^2^). In DM1-2 tissues, CAL led to only a small, non-significant increase (∼3.03 to ∼3.10–3.44 mN/mm^2^), and DM1-3 tissues remained weak, with CAL increasing force only modestly (∼0.39 to ∼0.65 mN/mm^2^).

Notably, despite the lack of improvement in peak tetanic force, CAL alleviated the fatigue observed in DM1-3 tissues during repeated tetanic stimulation (Supplementary Fig. 14a-c), indicating improved resistance to activity-induced decline. This functional benefit was accompanied by an increase in myotube CSA, most prominently in DM1-3 tissues, where CAL elicited the largest hypertrophic shift (Supplementary Fig. 14d-f).

We next examined whether calcitriol influenced fiber-type composition. When fiber types were pooled across patient lines, CAL appeared to shift the overall distribution toward a more control-like profile, with fewer slow type I fibers and pathological I/IIx hybrids, and a relative increase in fast type IIx and physiological IIa/IIx hybrids (Fig. 6e). However, donor-resolved analyses revealed modest and heterogeneous responses across DM1 subtypes (Fig. 6f–i). In DM1-1 tissues, CAL produced the clearest remodeling effect, increasing type IIx fibers from ∼12% to ∼38% and reducing pathological I/IIx hybrids from ∼70% to ∼45%, alongside a downward trend in type I fibers (from ∼12% to ∼3–4%, p = 0.08). DM1-2 tissues exhibited a mixed pattern: type I fibers decreased substantially (from ∼53% to ∼19%), whereas I/IIx hybrids increased (from ∼35% to ∼58%), accompanied by small increases in IIa/IIx hybrids (∼7% to ∼14%) and a non-significant trend toward higher IIx abundance (∼1–2% to ∼12%, p = 0.06). We observed no significant changes in DM1-3 tissues, although mean values suggested subtle shifts toward higher IIx (∼11% to ∼20%) and IIa/IIx (∼15% to ∼27%) proportions and lower type I content (∼28% to ∼12%). Overall, calcitriol induced limited, donor-dependent remodeling of fiber-type composition, underscoring the importance of accounting for DM1 heterogeneity in these analyses.

### Calcitriol alleviates myotonia despite persistent *CLCN1* splicing defects

Calcitriol induced coordinated regulation of ion channel and calcium-handling genes consistent with stabilization of muscle excitability. Among these changes, CAL significantly upregulated the calcium-selective channel *TRPV6*, a canonical vitamin D receptor target, together with stabilizing potassium channels *KCNK3* and *KCNQ5*, suggesting the establishment of a new conductance balance that counteracts hyperexcitability.

In parallel, genes involved in intracellular calcium handling, including the store-operated calcium entry regulators *CRACR2A* and *CRACR2B*, were upregulated. Conversely, CAL suppressed expression of excitability-promoting channels, including the sodium channel *SCN9A*, the calcium-activated chloride channel *ANO1 (TMEM16A)*, and the potassium channel *KCNN3*, as well as additional channel modulators, consistent with dampened membrane excitability. Collectively, these transcriptional changes point to activation of ionic compensation networks capable of offsetting the functional consequences of persistent *CLCN1* splicing defects (Fig. 6b).

Consistent with this molecular profile, CAL robustly rescued the myotonia-like phenotype across all DM1 subtypes (Fig. 6j,l). In juvenile DM1-1 tissues, the myotonia AUC decreased from ∼14.7 to ∼8.1, while DM1-2 tissues showed a similar reduction (∼8.6 to ∼5.0). The greatest improvement was observed in DM1-3 tissues, where AUC decreased from ∼3.5 to ∼1.7. These reductions were accompanied by near-complete normalization of relaxation between tetanic pulses, with force decay during resting phases resembling that of control tissues (Fig. 6k,l). Importantly, this functional rescue occurred despite persistent *CLCN1* mis-splicing (Supplementary Fig. 15), demonstrating that muscle hyperexcitability can be mitigated independently of chloride channel splice correction. Together, these findings highlight the utility of our 3D model in capturing functional disease phenotypes and revealing therapeutic benefits that are independent of MBNL1 upregulation.

## Discussion and conclusions

DM1 has lacked human *in vitro* models that directly integrate its defining molecular pathology with quantitative measurements of mature skeletal muscle function. Here, we establish a patient-derived 3D skeletal muscle platform that bridges this gap by simultaneously reproducing molecular, structural, and functional hallmarks of DM1 in a human context. Engineered tissues recapitulate RNA toxicity features—including nuclear RNA foci, MBNL1 sequestration, and widespread mis-splicing— while enabling expression and pathogenic mis-splicing of *CLCN1*. Functionally, the model captures impaired calcium handling, weakness, fatigue, and, critically, myotonia-like delayed relaxation, alongside disease-associated shifts in fiber-type composition. Together, these features position the platform as a robust system for mechanistic investigation and therapeutic testing across DM1 clinical subtypes.

Compared with existing DM1 models, this system uniquely links human RNA toxicity to directly measurable muscle physiology. Two-dimensional patient-derived cultures reliably reproduce nuclear RNA foci and selected splicing defects but fail to achieve the functional maturation required for contractile output or *CLCN1* expression, precluding analysis of myotonia *in vitro*^*70*^. Animal models, while instrumental in establishing causal links between spliceopathy and excitability defects, rely on supraphysiological repeat lengths and species-specific regulatory programs that complicate translation to human muscle. The emergence of *CLCN1* expression and pathogenic mis-splicing within the engineered 3D environment underscores the requirement for tissue maturation to unlock disease-relevant ion-channel regulation^71^, positioning this model as a necessary intermediate between 2D *in vitro* systems and *in vivo* studies.

Across patient subtypes, molecular pathology aligned with functional impairment, revealing shared disease mechanisms alongside clear donor-specific heterogeneity. RNA foci burden and MBNL1 sequestration tracked with CTG repeat length and overall functional severity, supporting a direct relationship between RNA toxicity and muscle dysfunction. Widespread spliceopathy affecting excitation–contraction coupling, membrane organization, sarcomeric integrity, and metabolic regulation provides a coherent molecular basis for the impaired calcium handling, weakness, and fatigability observed in the tissues. Notably, late-onset DM1-2 tissues retained near-normal maximal force yet displayed pronounced fatigability, demonstrating that endurance deficits can arise independently of baseline strength. This dissociation mirrors clinical observations in which fatigue progression does not strictly parallel muscle weakness ^*72*^, and highlights the importance of functional readouts beyond peak force.

In parallel, DM1 tissues exhibited a disease-associated shift in fiber-type distribution, characterized by enrichment of slow type I fibers and pathological hybrid phenotypes at the expense of fast glycolytic populations. Such fiber-type dysregulation, previously reported in patient muscle^52,73^ reflects a broader metabolic and contractile reprogramming rather than a simple atrophic response. Together, these findings indicate that DM1 muscle dysfunction arises from coordinated remodeling across molecular, structural, and functional levels, rather than from isolated defects in individual pathways.

Pharmacological treatment targeting RNA toxicity pathways revealed a critical distinction between molecular engagement and functional recovery in DM1 muscle. Although phenylbutazone reduced RNA foci and MBNL1 sequestration in the most affected subtypes, this partial molecular correction did not translate into restoration of splicing fidelity or improvement of calcium handling, force generation, fatigue resistance, or myotonia-like relaxation. In contrast, calcitriol elicited substantial functional improvements without altering MBNL1 localization or *CLCN1* splicing. This dissociation demonstrates that key aspects of DM1 muscle dysfunction—particularly excitability, fatigue, and relaxation—remain modifiable through mechanisms that operate downstream of or in parallel with RNA toxicity.

Importantly, functional rescue was most evident in the normalization of myotonia-like delayed relaxation, showing that membrane hyperexcitability in human DM1 muscle can be mitigated despite persistent *CLCN1* mis-splicing. This finding challenges the classical view that myotonia in DM1 is obligatorily dependent on restoration of chloride channel splicing and instead supports a model in which muscle excitability can be therapeutically stabilized through alternative ionic and signaling pathways. Such functional plasticity expands the mechanistic framework of DM1 myotonia and opens therapeutic opportunities beyond splice-correcting strategies.

Notably, CAL did not induce MBNL1 upregulation in our human 3D tissues, in contrast to reports in murine DM1 models^67^. Instead, the compensatory mechanisms engaged by CAL in human muscle operate independently of MBNL1 restoration. Together, these observations highlight the importance of human-specific tissue models for revealing therapeutic mechanisms that may be obscured or misrepresented in rodent systems.

At a broader level, CAL-associated functional rescue was accompanied by coordinated modulation of pathways supporting muscle performance, including excitation–contraction coupling, metabolic support, extracellular matrix organization, and ion handling. These adaptations provide a systems-level context for the observed improvements in calcium dynamics, fatigue resistance, and relaxation kinetics. While calcium handling improved across all DM1 subtypes, structural and metabolic adaptations were most evident in the most severely affected tissues, indicating subtype-dependent engagement of compensatory mechanisms. This heterogeneity underscores both the plasticity and the limits of functional rescue in DM1 muscle.

Beyond ion handling, CAL was also associated with attenuation of extracellular matrix and inflammatory transcriptional programs aberrantly activated in DM1 muscle. These changes are consistent with reduced profibrotic and stress-associated signaling and may contribute to improved tissue homeostasis and functional stabilization in the engineered muscles^74–76^.

Taken together, these findings define a therapeutic principle in which functional improvement in DM1 muscle can be achieved through coordinated activation of compensatory molecular networks rather than direct correction of the primary spliceopathy. In this framework, CAL does not restore a normal transcriptomic state but instead stabilizes muscle performance by engaging downstream pathways that offset the functional consequences of persistent RNA toxicity. This perspective positions compensatory network activation as a viable strategy for improving muscle function in DM1, particularly in settings where splice-correcting approaches alone may be insufficient.

Calcitriol also exerted modest and donor-dependent effects on fiber-type composition. While pooled analyses suggested partial shifts toward a more control-like distribution, donor-resolved responses were heterogeneous and insufficient to consistently restore the pathological fiber-type profile. These observations reinforce the importance of patient-specific context when evaluating therapeutic responses and underscore the utility of this platform for stratified analysis.

In conclusion, we present a functionally mature, patient-derived 3D human skeletal muscle model that integrates molecular fidelity with quantitative functional phenotyping to advance DM1 research. This is the first in vitro system to recapitulate both *CLCN1* mis-splicing and myotonia-like relaxation defects in human muscle while revealing therapeutic responses uncoupled from correction of MBNL1 localization or spliceopathy. By enabling mechanistic dissection and drug testing in a human-relevant context across DM1 subtypes, this platform provides a powerful foundation for preclinical discovery and the development of combinatorial therapeutic strategies.

## Methods

### Cell culture

Immortalized myoblasts from the biceps of three DM1 patients (DM1-1, DM1-2, and DM1-3), each with distinct clinical features, were previously established and characterized ^38^. Control myoblasts were derived from the quadriceps (CNT-1, CNT-2) and paravertebral muscle (CNT-3) of healthy donors. Cells were cultured at 37 °C in 5% CO_2_ and expanded in growth medium (GM) consisting of skeletal muscle basal medium supplemented with skeletal muscle supplement mix (PromoCell), 10% (v/v) fetal bovine serum (Gibco), and 100 U ml^-1^ penicillin with 100 µg ml^-1^ streptomycin (Gibco).

Differentiation medium (DM) was based on Dulbecco’s Modified Eagle Medium, high glucose, GlutaMAX™ Supplement (Gibco), containing 1% (v/v) KnockOut™ Serum replacement (Gibco), 1% (v/v) Insulin-Transferrin-Selenium-Ethanolamine (Gibco), 1% (v/v) Penicillin-Streptomycin-Glutamine (Gibco), 100 ng ml^-1^ human agrin (Sigma-Aldrich), 50 ng ml^-1^ human Insulin-like Growth Factor-1 (IGF-1, Sigma-Aldrich), and 1 mg ml^-1^ 6-aminocaproic acid (Sigma-Aldrich).

### Fabrication of DM1 patient-derived 3D skeletal muscle tissues

Silicon casting molds were fabricated via replica molding from custom 3D-printed molds, as described previously^30,32,77^. Each mold consisted of a rectangular well with a 30 µl capacity and two T-shaped flexible posts. Molds were sterilized under UV light and coated with 2% (w/v) Pluronic® F-127 (Sigma-Aldrich) for 2-24 h at 4 ºC.

Myoblasts were encapsulated by mixing a cell suspension (2.5 x 10^7^ cells ml^-1^) with a fibrin-based hydrogel and pipetting the mixture into the casting wells. The final hydrogel formulation contained 4 mg ml^-1^ fibrinogen from human plasma (Sigma-Aldrich), 2 U ml^-1^ thrombin from human plasma (Sigma-Aldrich), and 30% (v/v) Matrigel® Growth Factor Reduced Matrix (Corning). Cell-laden hydrogels were incubated at 37 ºC and 5% CO_2_ in GM containing 1 mg ml^-1^ 6-aminocaproic acid (Sigma-Aldrich). After 48 h, the media was replaced with DM, and half the volume was exchanged every 2 days.

### Treatment with small molecules

On day 13 of differentiation, DM1-derived tissues were treated with small molecules previously reported to upregulate MBNL1: phenylbutazone (PBZ, 324 µM; MedChemExpress), calcitriol (CAL, 1 µM; Selleckchem), Vorinostat (10 µM; Bio-Techne), and chloroquine diphosphate (10 µM; Sigma-Aldrich). Chloroquine was dissolved in water, whereas PBZ, CAL, and vorinostat were prepared in dimethyl sulfoxide (DMSO, ATCC) before dilution into the differentiation medium (DM). Because DMSO affects muscle contractility^30^, its final concentration was standardized at 0.01% (v/v) across all conditions, including untreated DM1 and healthy controls. Treatments were maintained for 96 h before endpoint assays.

### Electrical pulse stimulation assay

Bioengineered muscle tissues were subjected to electrical pulse stimulation (EPS) following 17 days of differentiation (including 4 days of treatment) to assess calcium dynamics and contractile function. Before stimulation, tissues were incubated with 30 µL of Fluo-8 dye-loading solution (Fluo-8 Calcium Assay Kit–No Wash, Abcam) for 1 h at 37 °C to enable detection of intracellular calcium flux. After incubation, the tissues were submerged in 1X phosphate-buffered saline (PBS) and transferred to a 24-well plate with 1 mL of fresh differentiation medium per well.

EPS was applied using Myo-MOVES, a custom-built stimulation setup consisting of graphite electrodes mounted on a 24-well plate lid^44^. The plate was placed on a Zeiss Axio Observer Z1/7 microscope equipped with the XL S1 cell incubator, maintaining samples at 37 °C and 5% CO_2_. Electrodes were connected to the Myo-MOVES well selector and a multifunction waveform generator (NF Corporation). Muscle tissues received a stimulation protocol of 15 periods of tetanic stimulation using a square stimulation signal (150 Hz, 2 s duration, 2 V mm^-1^ field strength, 10% duty cycle), with 5 s rest intervals between pulses.

### Immunohistochemistry and fluorescent methods

Tissues were fixed after 17 days of differentiation (Supplementary Fig. 1a) in 10% formalin solution (4% paraformaldehyde; Sigma-Aldrich), cryoprotected in 30% (w/v) sucrose in PBS, and embedded in OCT compound (Polyfreeze; Sigma-Aldrich), as previously described^30^. Transverse cryosections (20 µm) were prepared using a Leica CM1900 cryostat, mounted on SuperFrost Plus™ slides (Fisher Scientific), and stored at −20 °C until use.

RNA foci were detected by fluorescence in situ hybridization (FISH). Sections were incubated in prehybridization buffer (2× saline sodium citrate [SSC], 30% deionized formamide) for 30 min at RT and hybridized 2 h at 37 °C in a wet chamber with a Cy3-(CAG)_7_-Cy3-labelled probe (1:100) diluted in hybridization buffer (40% deionized formamide, 2×SSC, 100mg/mL dextran sulfate, 0.2% BSA, 2 mM ribonucleoside vanadyl complex (Sigma-Aldrich), 1 mg ml^-1^ E. coli tRNA, and 1 mg ml^-1^ herring sperm DNA. Samples were washed twice in prehybridization buffer at 42 °C (10 min each), followed by a PBS rinse.

For MBNL1 detection after FISH, sections were permeabilized 3 × 5 min in PBS-T (0.1% Triton X-100 in PBS), blocked in UltraCruz® Blocking Reagent (Santa Cruz Biotechnology) for 1 h, and incubated overnight at 4 °C with monoclonal anti-MBNL1 antibody (Supplementary Table 2), followed by signal amplification using a biotin-conjugated secondary antibody (Supplementary Table 3) and the VECTASTAIN® Elite® ABC Kit (Vector Laboratories) for 1 h at RT. After 3x PBS-T washes (5 min/wash), slides were incubated with Alexa Fluor™ 488 (1:200, Invitrogen) for 2 h and counterstained with VECTASHIELD® Antifade Mounting Medium with DAPI (Vector).

For myotube structural analysis, whole tissues or cryosections were permeabilized in PBS-T for 1 h, blocked in UltraCruz® Blocking Reagent (Santa Cruz Biotechnology) for 2 h at RT, and incubated overnight at 4 °C with primary antibodies (Supplementary Table 2). After PBS-T washes, samples were incubated for 2 h at RT with fluorophore-conjugated secondary antibodies (Supplementary Table 3) and counterstained with VECTASHIELD® Antifade Mounting Medium with DAPI (Vector).

To assess fiber-type composition, cryosections were incubated for 30 min at RT in blocking buffer (5% BSA, 1% normal goat serum in PBS), then for 1 h at 37 °C with a primary antibody cocktail targeting different Myosin Heavy Chain isoforms and dystrophin (Supplementary Table 2, diluted in 1% BSA in PBS. After PBS washing, sections were incubated for 1 h at 37 °C with the corresponding fluorophore-conjugated secondary antibodies (Supplementary Table 3). All incubations were performed in a wet chamber. Slides were mounted with Fluoromount-G™ (Thermo Fisher Scientific).

For immunofluorescence staining of CLCN1 and dystrophin, after fixation and cryopreservation as previously described, the tissues were transversely sectioned at a thickness of 10 µm. The protocol used for CLCN1 staining has been previously described. Briefly, tissue slides were permeabilized in 2% Acetone in 1× PBS (chilled) for 5 min at RT, washed 3 × 5 min in 1× PBS at RT, blocked in blocking buffer (1× PBS, 5% goat serum, 0.3% Triton X-100) containing 0.3 M glycine for 1 h at RT and incubated overnight at 4 °C with primary antibodies for CLCN1 and Dystrophin (Supplementary Table 2). Then, 3 × 5 min washes in 1× PBS at RT, slides were incubated for 1 h with secondary antibodies diluted in blocking buffer at RT (goat Alexa568 anti-rabbit IgG antibody (1:150) and goat Alexa488 anti-mouse IgG antibody (1:200, Supplementary Table 3). After washing 3 × 5 min in 1× PBS at RT, slides were mounted and counterstained with VECTASHIELD® Antifade Mounting Medium with DAPI (Vector).

### Imaging and image analysis

Brightfield images were acquired using a ZEISS Axio Observer Z1/7 microscope, and fluorescence images were obtained with a ZEISS LSM 800 confocal laser scanning microscope. Image analysis was performed using Fiji (ImageJ) software^78^.

For RNA foci and MBNL1 quantification, images were acquired using a ZEISS LSM 800 confocal microscope at 40× magnification and analyzed with ZEN Blue software and ImageJ. A total of 12 to 19 images per experimental condition were evaluated. The analysis procedure involved: (1) channel separation to select the red channel; (2) conversion to 8-bit format, followed by filtering for particle definition enhancement and background noise subtraction; and (3) automatic thresholding for particle segmentation. Using Fiji’s Analyze Particles function, the following parameters were quantified: number of nuclei, RNA foci per nucleus, MBNL1 aggregates per nucleus, RNA foci with MBNL1 colocalization per nucleus, and Corrected Total Cell Fluorescence (CTCF) for MBNL1 (AU/µm^2^). CTCF was calculated as CTCF = Integrated Density - (Area of selected cell × Mean fluorescence of background readings), providing a normalized measure of MBNL1 protein abundance that accounts for imaging variations and cell size differences.

For myotube size quantification, full transverse-section confocal images (20X magnification, tiled acquisition) of dystrophin-stained cryosections were segmented using Cellpose 2^79^. Individual cross-sectional areas (CSA) were then measured using the LabelsToRois plugin^80^ in Fiji. Across samples, segmentation was robust and typically identified 1200-1900 myotubes per section. A minimum of five independent tissue samples were analyzed per condition. The effective CSA of each sample was calculated by summing the CSA of all myotubes within each image.

Confocal images of MYH-stained cryosections were processed for muscle fiber-type quantification using the same segmentation workflow based on dystrophin labeling. ROIs generated with LabelsToRois were overlaid onto the corresponding MYH isoform channels. Then, mean gray intensity values were measured to determine individual fiber types and quantify overall sample composition. A minimum of three independent tissue samples were analyzed per condition.

### Video processing and force measurements

Brightfield and fluorescence videos during EPS were acquired using the ZEISS Axio Observer Z1/7 microscope and analyzed with Fiji (Image J). For calcium transient analysis, fluorescent videos of the entire top view of the tissues were obtained, and the mean gray value per frame was measured. Changes in fluorescence were then normalized to the baseline signal (ΔF/F0).

Contractile force generated by 3D skeletal muscle tissues in response to EPS was quantified by measuring the lateral displacement of the flexible PDMS posts, as previously described^30,81^. Displacement was analyzed in ImageJ, and force was calculated using a custom Python script based on Euler–Bernoulli beam theory:

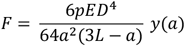

Where *F* is the applied force, *E* the Young’s modulus of PDMS (1.6 MPa), *D* the post diameter, *a* the height at which force is applied, *L* the recording height of the post, and *y(a)* the observed displacement. Force values were normalized to the effective CSA of each tissue (see previous section) to compute specific force^82^.

### Western blotting

For total protein extraction, 3D muscle tissues were homogenized using a TissueLyser (Qiagen) in RIPA buffer (Fisher Scientific) supplemented with protease and phosphatase inhibitor cocktails (Roche Applied Science). Total protein content was quantified using a BCA assay kit (Thermo Scientific Pierce) following the manufacturer’s instructions. For specific protein detection, 10 µg of total protein were denatured for 5 min at 100°C, electrophoresed on 12% SDS-PAGE gels, and transferred to 0.45 µm PVDF membranes using wet transfer for 1.5 h at 23V. Before transfer, PVDF membranes were pre-activated by immersing them in methanol for 30 seconds, followed by equilibration in the transfer buffer. Membranes were blocked with EveryBlot Blocking Buffer (Bio-Rad) for 1 h at room temperature with gentle agitation. Membranes were incubated overnight at 4°C with blocking solution containing mouse anti-MBNL1 (1:200, clone MB1a, Developmental Studies Hybridoma Bank).

For MBNL1 detection, membranes were subsequently incubated with goat anti-Mouse IgG (H+L) Cross-Adsorbed Secondary Antibody, HRP secondary antibody (1:3000, Thermo Fisher Scientific) for 1 h at room temperature. Following detection, membranes were stripped using mild stripping buffer (0.2 M NaOH with 0.1% SDS) for 15 minutes at room temperature with gentle agitation. Membranes were then washed 3 times for 5 minutes each in TBST (Tris-buffered saline with 0.1 % Tween 20), reblocked with EveryBlot Blocking Buffer (Bio-Rad) for 30 min at room temperature, and incubated for 2 h at room temperature with goat HRP-conjugated anti-GAPDH (1:800, Santa Cruz Biotechnology, clone G-9) as a loading control. Immunoreactive bands were visualized using Signal West Pico PLUS Chemiluminescent Substrate (Thermo Fisher Scientific), and images were acquired using an iBright™ FL1500 Imaging System (Invitrogen). Quantitative analysis of the images obtained was conducted using ImageJ (NIH) software.

### RNA isolation, cDNA synthesis, and Splicing analysis

Post-treatment tissues were flash frozen in liquid nitrogen. For RNA extraction, tissues were homogenized using TissueLyser LT (Qiagen) in QIAzol reagent (Qiagen), and the total RNA was extracted using the miRNeasy Micro Kit (Qiagen) according to the manufacturer’s instructions. RNA quantification was assessed using a Nanodrop ND-1000 full-spectrum spectrophotometer. 600 ng of RNA was retrotranscribed to cDNA using the QuantiTect Reverse Transcription Kit (Qiagen) according to the manufacturer’s protocol. Alternative splicing was analyzed using 25-50 ng of cDNA in a standard PCR reaction with GoTaq polymerase (Promega, Inc.) and specific primers. Primers and PCR conditions are described in Supplementary Table 4. PCR products were separated in a 2.5% agarose gel and quantified using ImageJ software (NIH).

### RNA-Seq mRNA library preparation, sequencing and analysis

RNA-seq libraries were prepared using the Illumina Stranded Total RNA Prep with Ribo-Zero Plus kit (Illumina), following the manufacturer’s instructions, starting from 0.5 µg of total RNA. Final libraries were assessed on an Agilent 2100 Bioanalyzer using the DNA 7500 assay for quality control. Libraries were sequenced on a NovaSeq 6000 system (Illumina) with paired-end 2×51 bp reads using the manufacturer’s dual-indexing protocol. Image analysis, base calling, and quality scoring were carried out with Real Time Analysis (RTA v3.4.4), and FASTQ files were generated with the corresponding Illumina pipeline.

RNA-seq analyses were performed in R (version 4.4.1, RFoundation for Statistical Computing). Reads were aligned to the human reference genome GRCh38 using STAR v2.7.8a^83^ with ENCODE-recommended parameters, and gene-level abundances were quantified with RSEM v1.3.0^84^ using GENCODE v44 annotation. Differential gene expression analysis was conducted with limma^85^ using the voom-transformed count data^86^, including appropriate normalization and statistical modeling steps. Multidimensional scaling plots were generated with the limma function plotMDS to visualize sample relationships. Functional enrichment analyses were performed separately using ShinyGO v0.82^87^, applying default settings unless otherwise specified, to identify significantly enriched biological pathways and gene ontology categories. For Vitamin D target gene selection and analysis, a predefined panel of established vitamin D– responsive genes was compiled from published literature^88^. Additional genes relevant to skeletal muscle biology (*VDR, MYOG*, and *CDKN1A/P21)* were included a priori based on their functional relevance. Genes were considered significantly regulated using an adjusted p-value < 0.05. Gene ontology enrichment analysis was performed using ShinyGO v0.82 with default statistical parameters, and network visualizations were generated using a similarity cutoff of 0.4.To evaluate transcriptional maturation of our 3D skeletal muscle tissues, we performed a comparative analysis using bulk RNA-seq data generated in this study together with publicly available datasets from Todorow et al.^62^, which include primary human myoblasts and myotubes derived from control and DM1 donors. For cross-platform comparison, raw count matrices were normalized and processed using the same pipeline as our samples.

To identify maturation-associated gene programs, we focused on transcripts with zero expression in 2D cultures but detectable expression in both human muscle biopsies and our 3D constructs. Gene ontology (GO) enrichment analysis was performed using ShinyGO v 0.82, applying default settings and an FDR threshold of <0.05. Additional enrichment analyses were performed using DAVID Bioinformatics Resources, with an EASE threshold of 0.08 and Benjamini–Hochberg FDR <0.05. DAVID identified significant overrepresentation of actin filament-based processes and contractile fiber categories among genes absent in 2D cultures but expressed in 3D constructs and biopsies. Heatmaps of selected genes were generated to illustrate representative members of these categories, including myosin heavy chain isoforms (*MYH1, MYH2, MYH3, MYH8)*, cytoskeletal components (*ACTA1, ACTA2)*, troponin proteins (*TNNI2)*, and sarcomeric structural proteins (*NEB, TTN, TPM4*).

Metabolic pathway enrichment in DAVID highlighted genes involved in oxidative phosphorylation, mitochondrial biogenesis, ATP synthesis, and aerobic metabolism, consistent with maturation-associated metabolic reprogramming in 3D tissues. All enrichment analyses were conducted using gene lists restricted to transcripts that met pre-defined expression thresholds and appropriate statistical significance criteria as described above.

## Statistical analysis

Statistical analyses were performed using GraphPad Prism. Data are presented as mean ± SEM. Comparisons between two groups were performed using a two-tailed Student’s t-test, with Welch’s correction applied when appropriate. Comparisons involving multiple groups were analyzed using one-way or two-way ANOVA, including repeated-measures two-way ANOVA where applicable, followed by Tukey’s multiple comparisons test.

Statistical significance was defined as P < 0.05 (*P< 0.05; **P < 0.01; ***P < 0.001; ****P< 0.0001). Sample sizes and the number of independent experiments are indicated in the figure legends.

## Acknowledgements

This work was supported by the Ministerio de Ciencia e Innovación (Spain) through Grant PID2022-139605OA-I00 (J.F.C.), and by the CIPROM/2023/22 project from the Generalitat Valenciana (R.A.). X.F.G. and M.S.A. were supported by postdoctoral fellowships from the Generalitat Valenciana (CIAPOS/2022/106 and CIAPOS/2022/122, respectively). The authors wish to acknowledge MyoLine, the platform for the immortalization of human cells at the Institute of Myology (Paris), and the Developmental Studies Hybridoma Bank, created by the NICHD of the NIH and maintained at The University of Iowa, for providing the monoclonal antibodies used in this study. The authors also thank the members of the Biosensors for Bioengineering Group (IBEC) for their valuable feedback and support.

## Author contributions

* These authors contributed equally: Xiomara Fernández-Garibay & María Sabater-Arcís. J.F.C., J.R.A., and R.A. conceived the study and developed the initial hypothesis. X.F.G., M.S.A. and J.F.C. designed the experimental approach. X.F.G. and M.S.A. performed all experiments. A.T.V. established the protocol for force measurement analysis. X.F.G., M.S.A., and J.F.C. analyzed and interpreted the data. K.M. and A.B. generated and provided the immortalized cell lines used in this work. M.S. and G.N.G. provided DM1 patient-derived cells and contributed to data interpretation and discussion. X.F.G. and M.S.A. wrote the manuscript with input from all authors. J.R.A. and J.F.C. supervised the study.

