## Supplementary info for "Patient-derived 3D engineered human muscle model recapitulates *CLCN1* mis-splicing and myotonia in myotonic dystrophy type 1"

### Supplementary Material

Supplementary Table 1. Immortalized myotonic dystrophy type 1 myoblasts from patients with genetic and clinical heterogeneity[1]

| Donor/Patient myoblasts | CNT-1 | CNT-2 | CNT-3 | DM1-1 | DM1-2 | DM1-3 |
|---|---|---|---|---|---|---|
| **Sex** | M | F | M | F | F | F |
| **Age at sampling** | 38 | 53 | 16 | 36 | 46 | 39 |
| **Age of onset** | - | - | - | 15 (juvenile) | 42 (late onset) | 27 (adult) |
| **Muscle involvement** | - | - | - | moderate | mild | moderate |
| **Cardiac involvement** | - | - | - | mild | mild | severe |
| **Respiratory involvement** | - | - | - | moderate | severe | mild |
| **mRS** | - | - | - | 2 | 1 | 4 |
| **Mode CTG expansion** | - | - | - | 953<br>2080 | 379<br>863 | 1224<br>2301 |
| **Type of muscle** | quadriceps | quadriceps | paravertebral | biceps | biceps | biceps |

Supplementary Table 2. Primary antibodies

| Class | Host | Target | Isotype | Supplier (Reference) | Technique (Dilution) |
|---|---|---|---|---|---|
| Monoclonal | Mouse | Muscleblind-like splicing regulator 1 1 protein (MBNL1) | IgG1 | Developmental Studies Hybridoma Bank (DSHB) (MB1a(A8)) | IF (1:100); WB (1:200) |
| Polyclonal | Rabbit | Sarcomeric a-actinin (SAA) | IgG | Abcam (ab137346) | IF (1:100) |
| Monoclonal | Mouse | Dystrophin (exons 31/32) | IgG2A | DSHB (MANDYS1(3B7)) | IF (1:50) |
| Monoclonal | Mouse | Dystrophin (exon 43) | IgG2A | DSHB (MANDYS106(2C6)) | IF (1:50) |
| Polyclonal | Rabbit | Dystrophin | IgG | Abcam (ab15277) | IF (1:200) |
| Monoclonal | Mouse | MYH1, Myosin heavy chain 1 Type IIX | IgM | DSHB (6H1) | IF (1:10) |
| Monoclonal | Mouse | MYH2, Myosin heavy chain Type IIA | IgG1 | DSHB (SC-71) | IF (1:100) |
| Monoclonal | Mouse | MYH7, Myosin heavy chain Type I | IgG2b | DSHB (BA-D5) | IF (1:30) |
| Polyclonal | Rabbit | Anti-CLCN1 antibody - C-terminal | IgG | ab189857 | IF (1:50) |
| Monoclonal | Mouse | GAPDH-HRP | IgG1 | sc-365062 HRP (G-9) | WB (1:800) |

Supplementary table 3. Secondary antibodies

| Class | Host | Reactivity / Target | Conjugate | Supplier (Reference) | Technique (Dilution) |
|---|---|---|---|---|---|
| Polyclonal | Goat | Rabbit/IgG | Alexa Fluor™ 488 | Invitrogen (A-11034) | IF (1:200) |
| Polyclonal | Goat | Mouse/IgG | Alexa Fluor™ Plus 647 | Invitrogen (A32728) | IF (1:200) |
| Polyclonal | Goat | Rabbit/IgG | Alexa Fluor™ 568 | Invitrogen (A-11036) | IF (1:200) |
| Polyclonal | Goat | Mouse/IgM | Alexa Fluor™ 488 | Invitrogen (A-21042) | IF (1:200) |
| Polyclonal | Goat | Mouse/IgG1 | Alexa Fluor™ 647 | Invitrogen (A-21240) | IF (1:200) |
| Polyclonal | Goat | Mouse/IgG2 | Alexa Fluor™ 350 | Invitrogen (A-21140) | IF (1:150) |
| Polyclonal | Goat | Mouse/IgG | HRP | ThermoFisher scientific (G-21040) | WB (1:3000) |

Supplementary Table 4. Primers and conditions used for RT-PCR splicing analysis.

| PCRs Transcript | Forward primer (5'→3') | Reverse primer (5'→3') | Annealing T° | Cycles | Fragment sizes (bp) | Reference |
|---|---|---|---|---|---|---|
| *BIN1* (exon 11) | AGAACCTCAATGATGTGCTGG | TCGTGTTGACTCTGATCTCGG | 58 | 28 | 163-208 | 2 |
| *MBNL1* (exon 7) | GCCCAATACCAGGTCAACCA | GGCCTCTTTGGTAATGGGGG | 58 | 35 | 101-155 | 3 |
| *LDB3* (exon 11) | GCAAGACCCTGATGAAGAAGCTC | GACAGAAGGCCGGATGCTG | 61 | 28 | 163-352 | 4 |
| *KIF13A* (exon 32) | TCCTGTCAGTATCCATCGGCT | TGAGTGCATCTGACCACCTCT | 65 | 30 | 117-156 | 1 |
| *ATP2A1* (exon 22) | CTCATGGTCCTCAAGATCTCAC | AGCTCTGCCTGAAGATGTGTCAC | 58 | 35 | 161-203 | 5 |
| *DMD* (exon 78) | TTAGAGGAGGTGATGGAGCA | GATACTAAGGACTCCATCGC | 58 | 28 | 116-148 | 6 |
| *INSR* (exon 11) | CCAAAGACAGACTCTCAGAT | AACATCGCCAAGGGACCTGC | 60 | 35 | 131-167 | 7 |
| *CLCN1* (exon 6b/7a) | CATCTCTCCCCAGGCTGT | GCATCCTTGTTCCACACT | 60 | 27 | 568-490-435 | 8 |
| *GAPDH* | CATCTTCCAGGAGCGAGATC | GTTCACACCCATGACGAACAT | 57 | 29 | - | 9 |

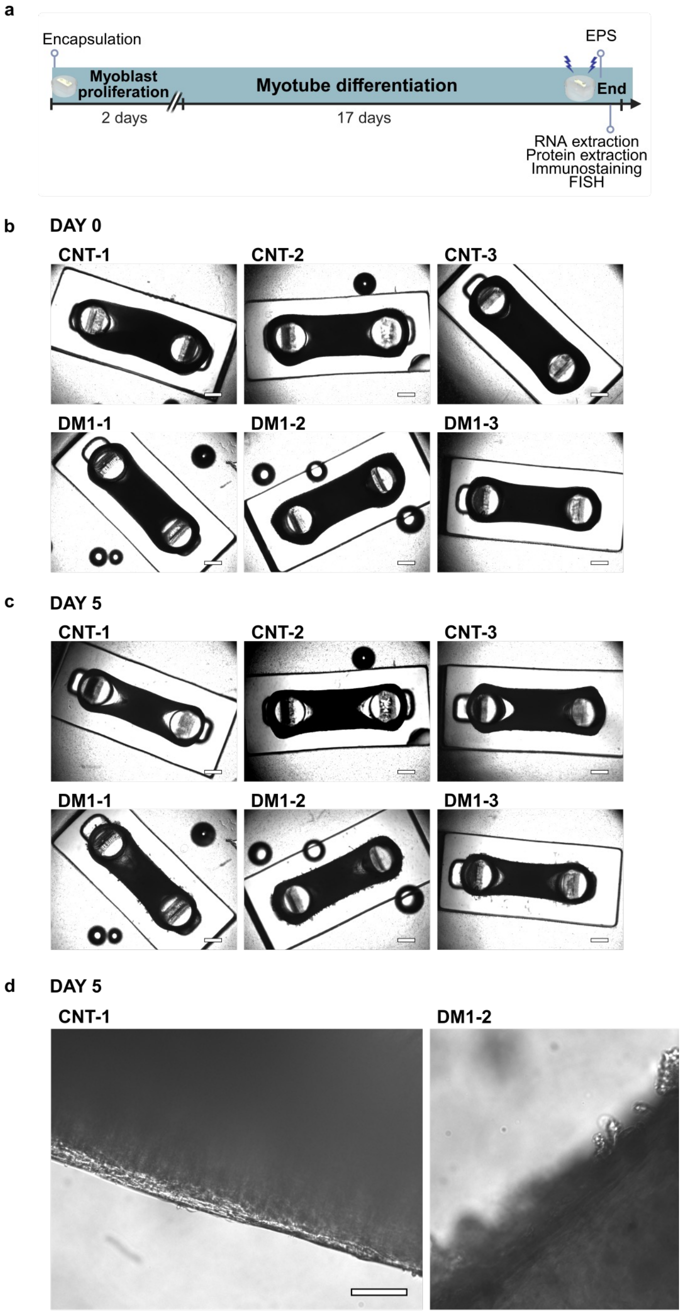


**Supplementary Fig. 1 | Experimental timeline and tissue compaction during 3D muscle differentiation.**
**a,** Schematic of the experimental workflow showing myoblast proliferation for 2 days followed by myotube differentiation for 17 days, with indicated endpoints for downstream analyses. **b–d,** Representative top-view brightfield images of healthy control (CNT) and DM1 3D muscle tissues illustrating tissue compaction over time. **(b)** Day 0. **(c)** Day 5. **(d)** Higher-magnification view at Day 5. Scale bars = 500 μm **(b,c)**; 100 μm **(d)**.

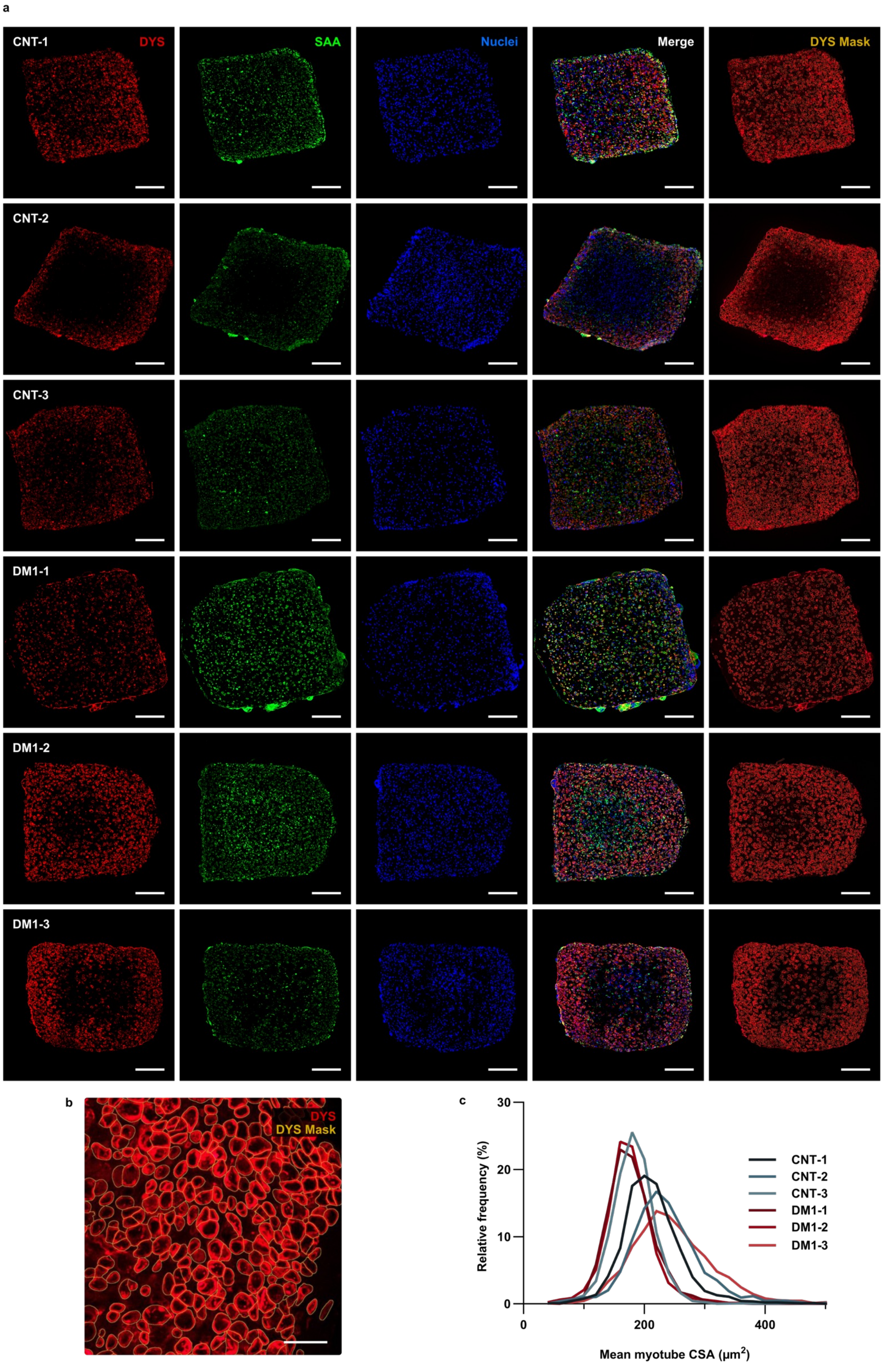
a
CNT-1
DYS
SAA
Nuclei
Merge
DYS Mask
CNT-2
CNT-3
DM1-1
DM1-2
DM1-3
b
DYS
DYS Mask
c
Relative frequency (%)
30
20
10
0
0
200
400
Mean myotube CSA (μm²)
CNT-1
CNT-2
CNT-3
DM1-1
DM1-2
DM1-3

**Supplementary Fig. 2 | Structural characterization and cross-sectional area analysis of CNT and DM1 3D muscle tissues.**
**a,** Representative confocal images of transversal cross-sections of CNT and DM1 tissues. Dystrophin (DYS, red), sarcomeric α-actinin (SAA, green), and nuclei (DAPI, blue) are shown, together with merged images. The calculated dystrophin mask (DYS mask) used for segmentation is overlaid as a yellow outline. Scale bars = 200 μm. **b,** Higher-magnification view showing DYS signal (red) and the corresponding DYS mask (yellow outline). Scale bar = 40 μm. **c,** Relative frequency distribution of myotube cross-sectional area (CSA) quantified from DYS mask images using Cellpose.

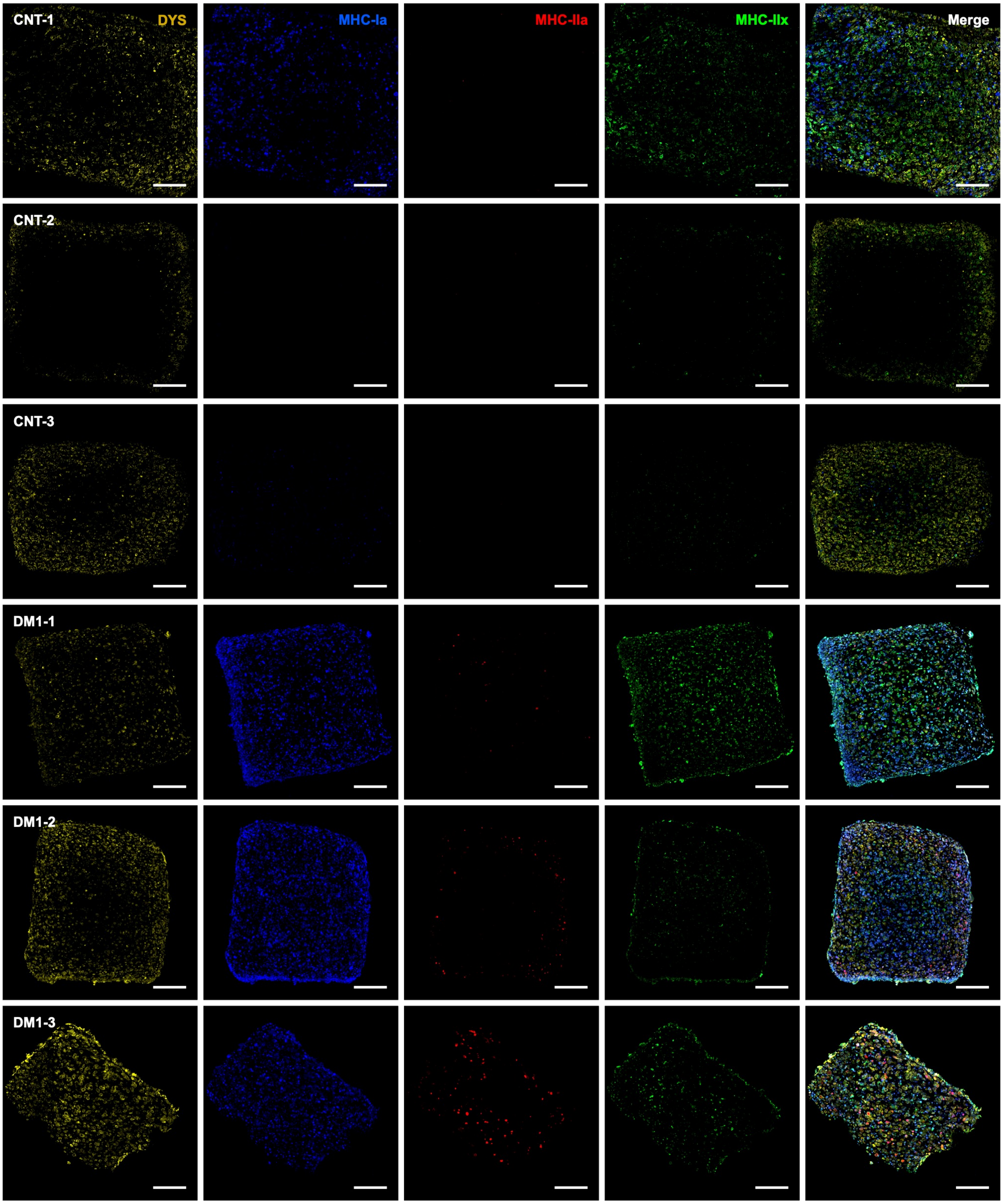


**Supplementary Fig. 3 | Fiber-type composition in CNT and DM1 3D muscle tissues.**
Representative immunofluorescence images of transversal cross-sections of CNT and DM1 tissues. Dystrophin (DYS, yellow) marks myofiber membranes, and myosin heavy chain (MyHC) isoforms identify fiber types: MyHC-I (blue), MyHC-IIx (green), and MyHC-IIa (red). Merged images show the distribution of MyHC isoforms within the 3D muscle tissues. Scale bars = 200 μm.

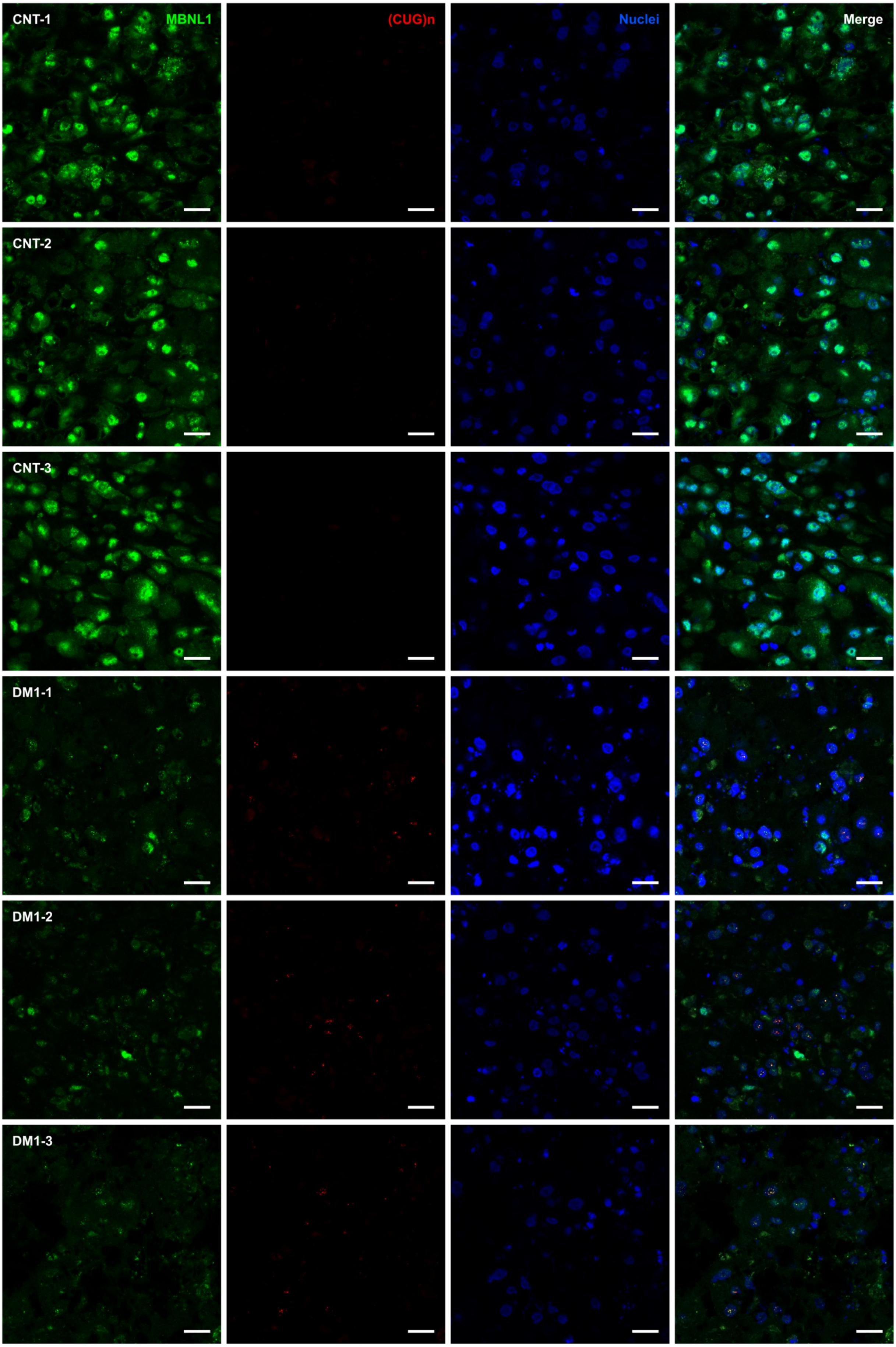


**Supplementary Fig. 4 | Ribonuclear RNA foci and MBNL1 localization in CNT and DM1 3D muscle tissues.**
Representative confocal images of transversal cross-sections of CNT and DM1 tissues. Ribonuclear (CUG)n RNA foci were detected by fluorescence *in situ* hybridization using a Cy3-labelled probe (red), MBNL1 was immunostained (green), and nuclei were counterstained with DAPI (blue). Scale bars = 20 μm.

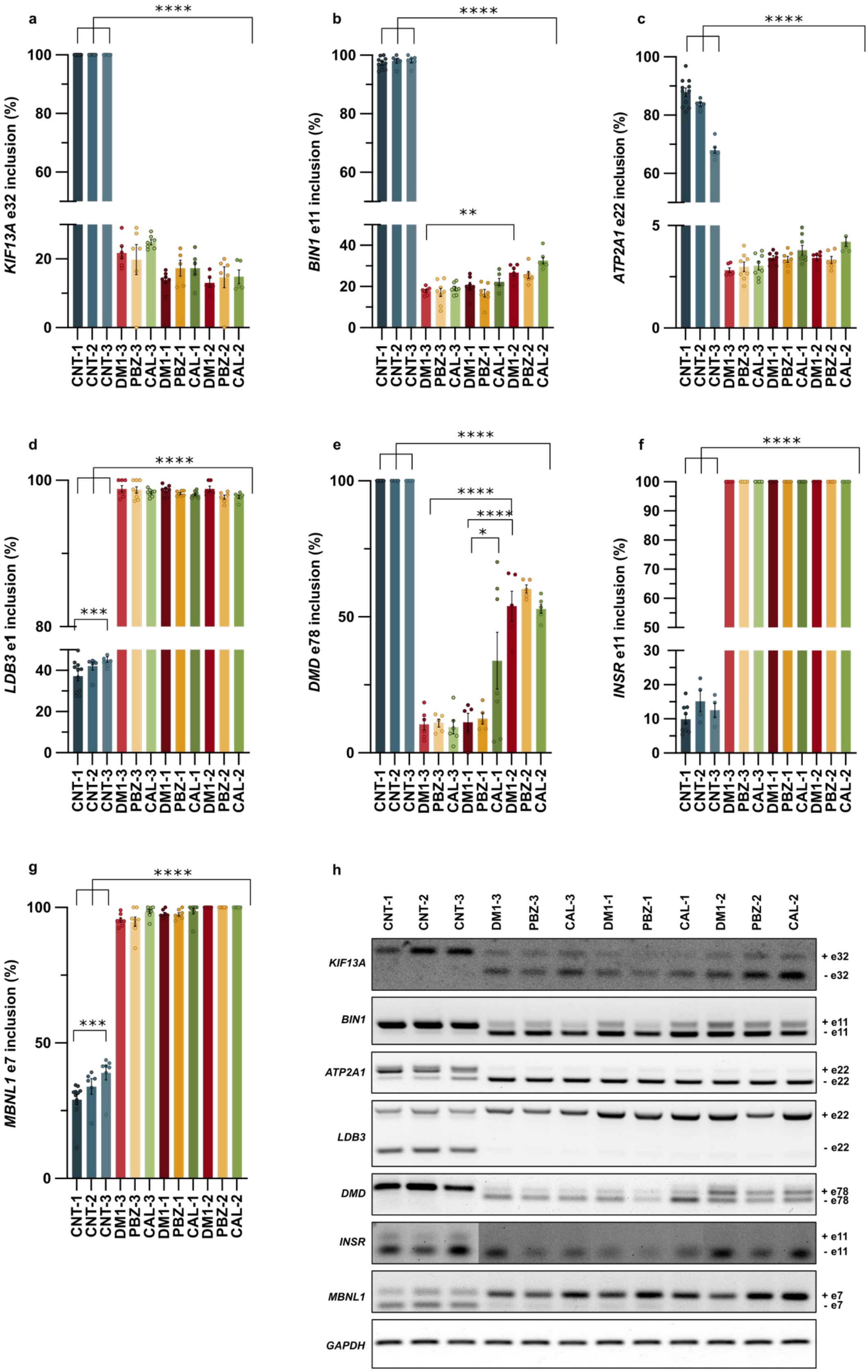


**Supplementary Fig. 5 | Alternative splicing defects are preserved in DM1 3D muscle tissues and are not corrected by PBZ or CAL treatment.**
**a–g,** Percentage of exon inclusion measured by semi-quantitative RT–PCR for **(a)** *KIF13A* (exon 32), **(b)** *BIN1* (exon 11), **(c)** *ATP2A1* (exon 22), **(d)** *LDB3* (exon 11), **(e)** *DMD* (exon 78), **(f)** *INSR* (exon 11), and **(g)** MBNL1 (exon 7) across healthy control (CNT), untreated DM1, and DM1 tissues treated with phenylbutazone (PBZ) or calcitriol (CAL). GAPDH was used as an internal control. **h,** Representative RT–PCR gels used for quantification in panels **(a–g)**. Sample size was n = 6–11 per condition. Bar graphs show mean ± SEM. Statistical significance was determined by one-way ANOVA followed by Tukey's HSD post hoc test. *P < 0.05; **P < 0.01; ***P < 0.001; ****P < 0.0001.

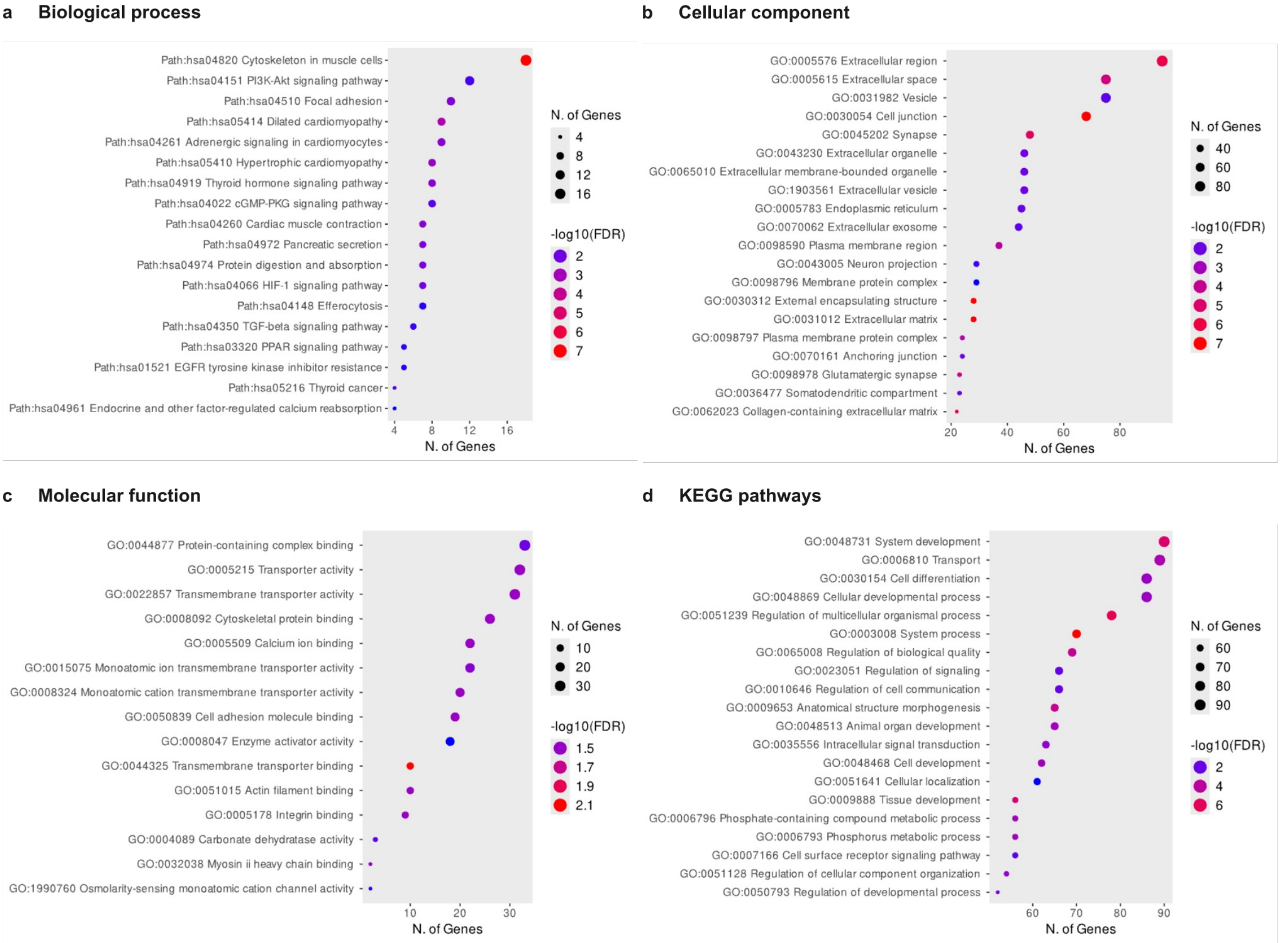


**Supplementary Fig. 6 | Transcriptomic differences between DM1 and control 3D muscle tissues.**
Gene Ontology (GO) and KEGG pathway enrichment analysis of differentially expressed genes between DM1 and healthy control (CNT) 3D muscle tissues. Enrichment results are shown for **(a)** biological process, **(b)** cellular component, **(c)** molecular function, and **(d)** KEGG signaling pathways. Dot size represents the number of genes associated with each term, and color intensity indicates statistical significance ($-\log_{10}$ FDR).

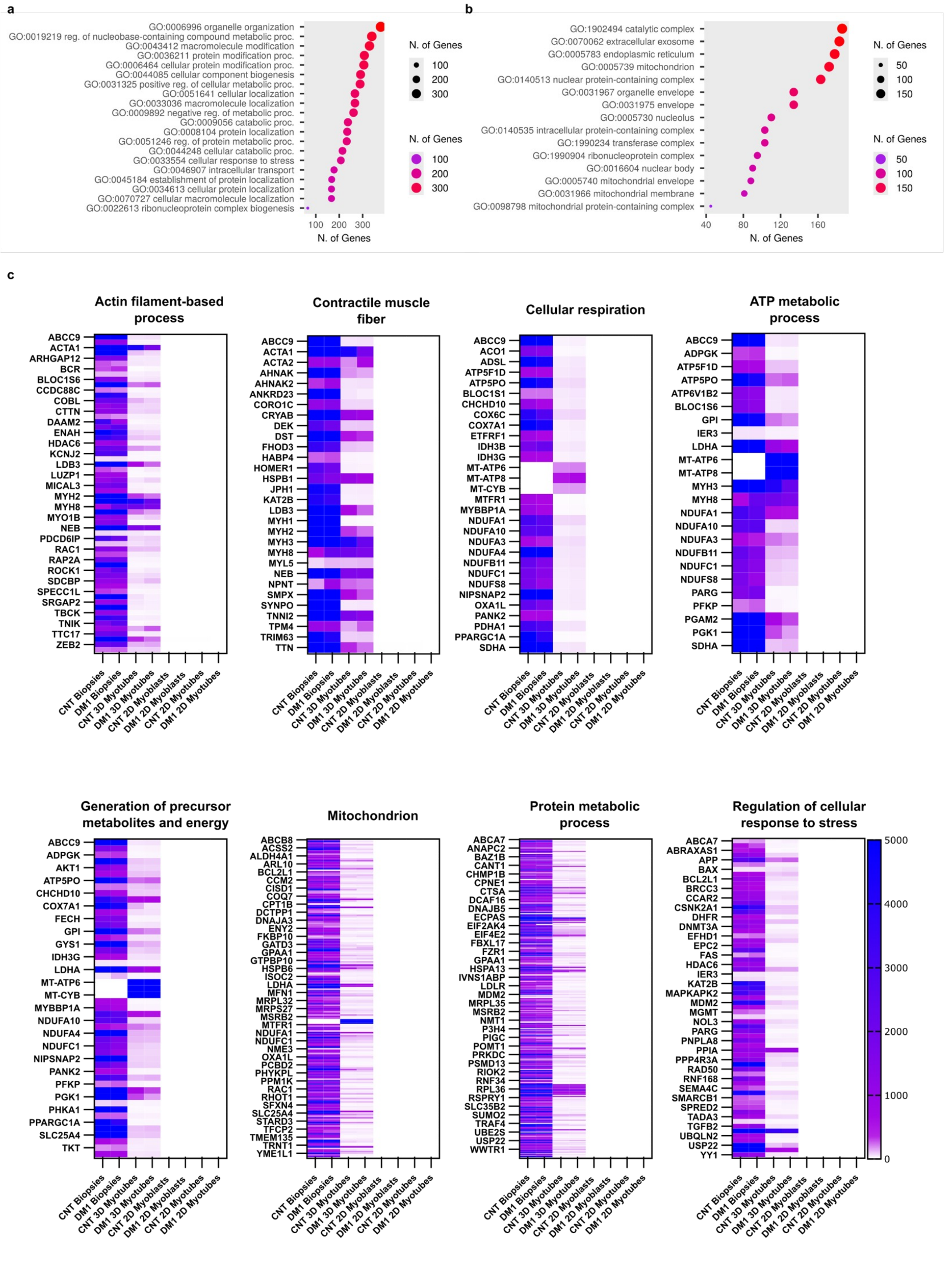


**Supplementary Fig. 7 | Transcriptomic comparison of 3D muscle tissues with 2D cultures and human muscle biopsies.**
Transcriptomic analysis of genes selectively expressed in 3D muscle tissues and human muscle biopsies but absent in 2D cultures. Genes with zero expression in 2D cultures (expression counts = 0) and detectable expression in both human muscle biopsies and 3D muscle tissues were selected using reference datasets from Todorow et al. (2021). **a,b,** Gene Ontology enrichment analysis of the selected gene set performed using ShinyGO (version 0.82). **c,** Functional annotation clustering of the same gene set using DAVID (EASE score threshold = 0.08; Benjamini–Hochberg FDR ≤ 0.05).

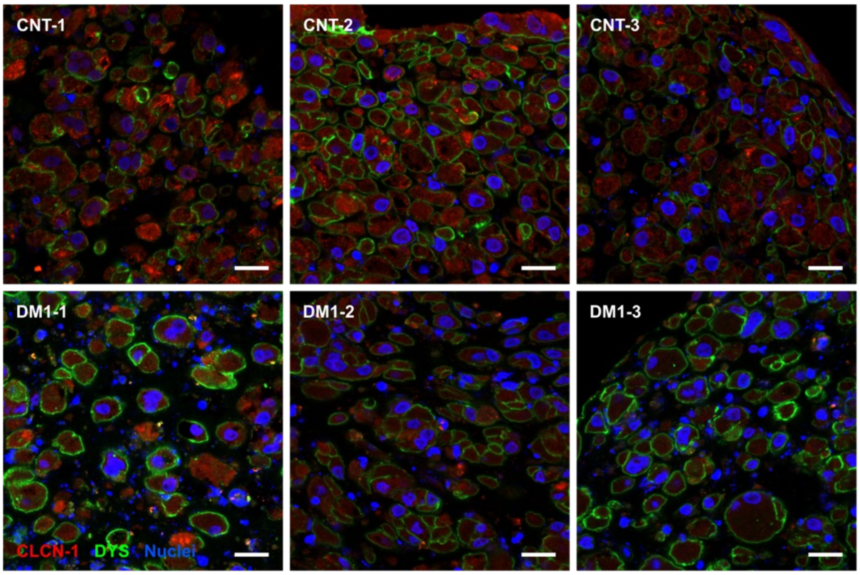


**Supplementary Fig. 8 | First detection of CLCN-1 in an *in vitro* DM1 model.**
Representative confocal images of transversal cross-sections showing CLCN-1 protein (red) in control (CNT) and DM1 3D muscle tissues. Myotube membranes were immunostained for dystrophin (DYS, green), and nuclei were counterstained with DAPI (blue). Scale bars = 20 μm.

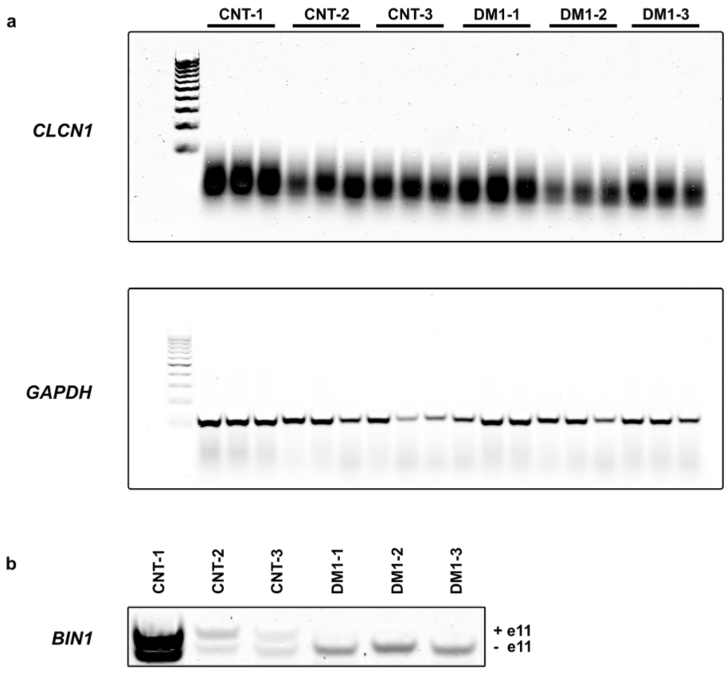


**Supplementary Fig. 9 | CNT and DM1 2D myotubes do not express *CLCN1*.**
**a,** Representative RT–PCR analysis of *CLCN1* expression in control (CNT) and DM1 2D myotubes, performed using the same conditions as for the corresponding 3D tissues from the same cell lines. The smear observed corresponds to primer dimers. *GAPDH* was used as an internal control. **b,** Representative RT–PCR gel showing *BIN1* exon 11 splicing alterations in CNT and DM1 2D myotubes, consistent with previously reported defects in these myoblast lines (Núñez-Manchón *et al.* 2024).

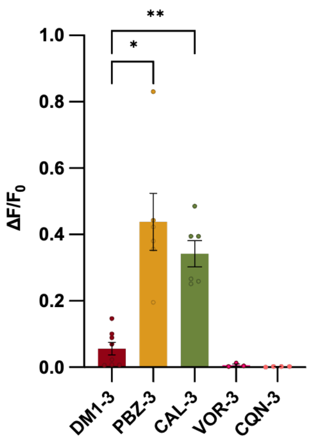


**Supplementary Fig. 10 | Effect of different small molecules on calcium transients in DM1 muscle tissues.**
Maximum calcium transient amplitude ($\Delta F/F_0$) measured during tetanic stimulation (150 Hz) in DM1-3 3D muscle tissues treated with phenylbutazone (PBZ, 324 µM), calcitriol (CAL, 1 µM), vorinostat (VOR, 10 µM), or chloroquine (CQN, 10 µM), compared with untreated DM1 control (DMSO 0.01%).
Data are shown as mean ± SEM. Statistical analysis was performed using one-way ANOVA followed by Dunnett's post hoc test comparing each treatment to untreated DM1 control. *$P < 0.05$; **$P < 0.01$. n = 3–6 tissues per group.

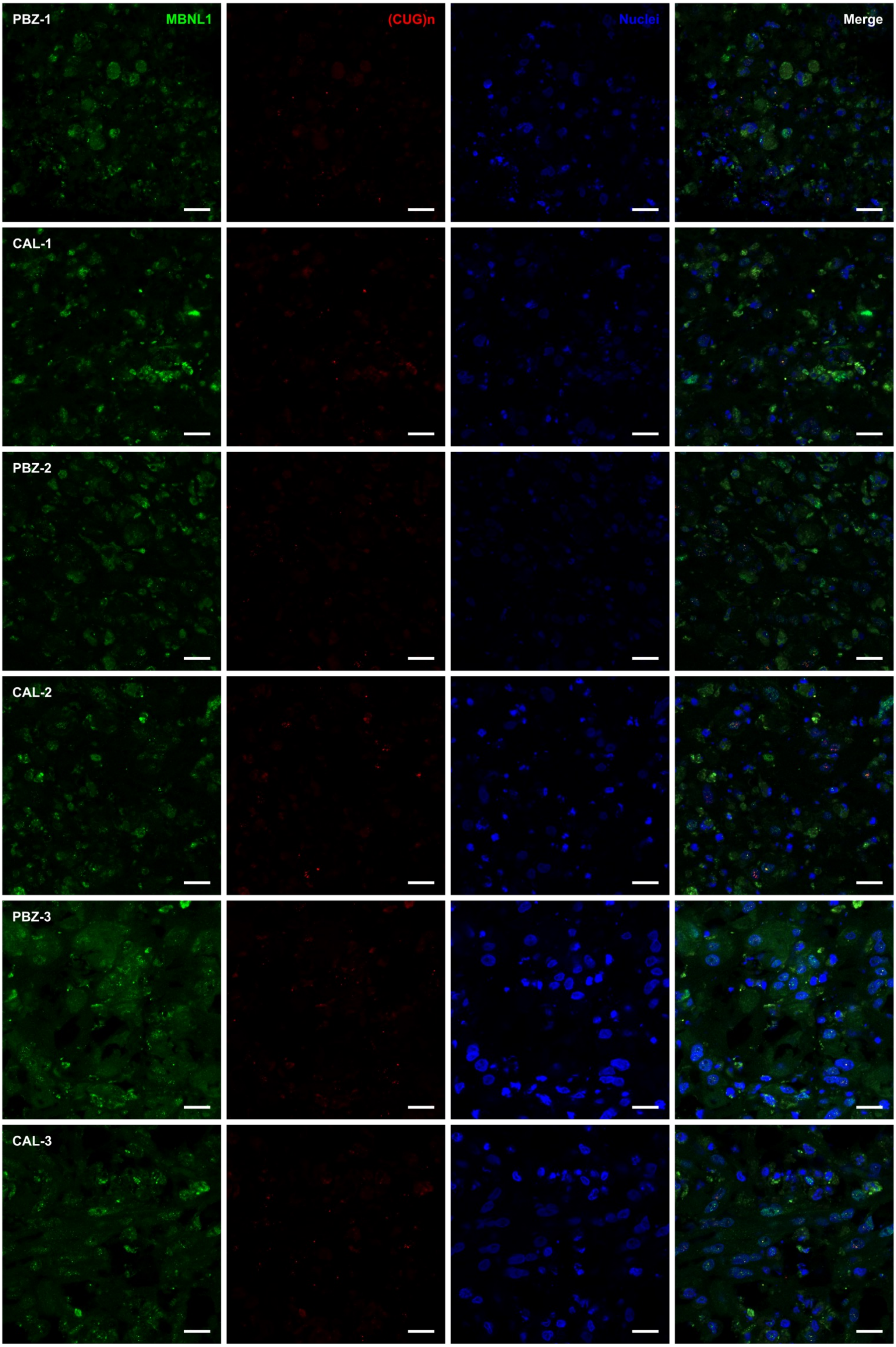


**Supplementary Fig. 11 | Representative confocal images of MBNL1 and (CUG)n staining in PBZ- and CAL-treated DM1 3D muscle tissues.**
Representative confocal images of transversal cross-sections showing MBNL1 immunostaining (green) and (CUG)n RNA foci detected by fluorescence *in situ* hybridization (red) in the three DM1 3D muscle tissue lines treated with phenylbutazone (PBZ) or calcitriol (CAL). Nuclei were counterstained with DAPI (blue). Scale bar = 20 μm.

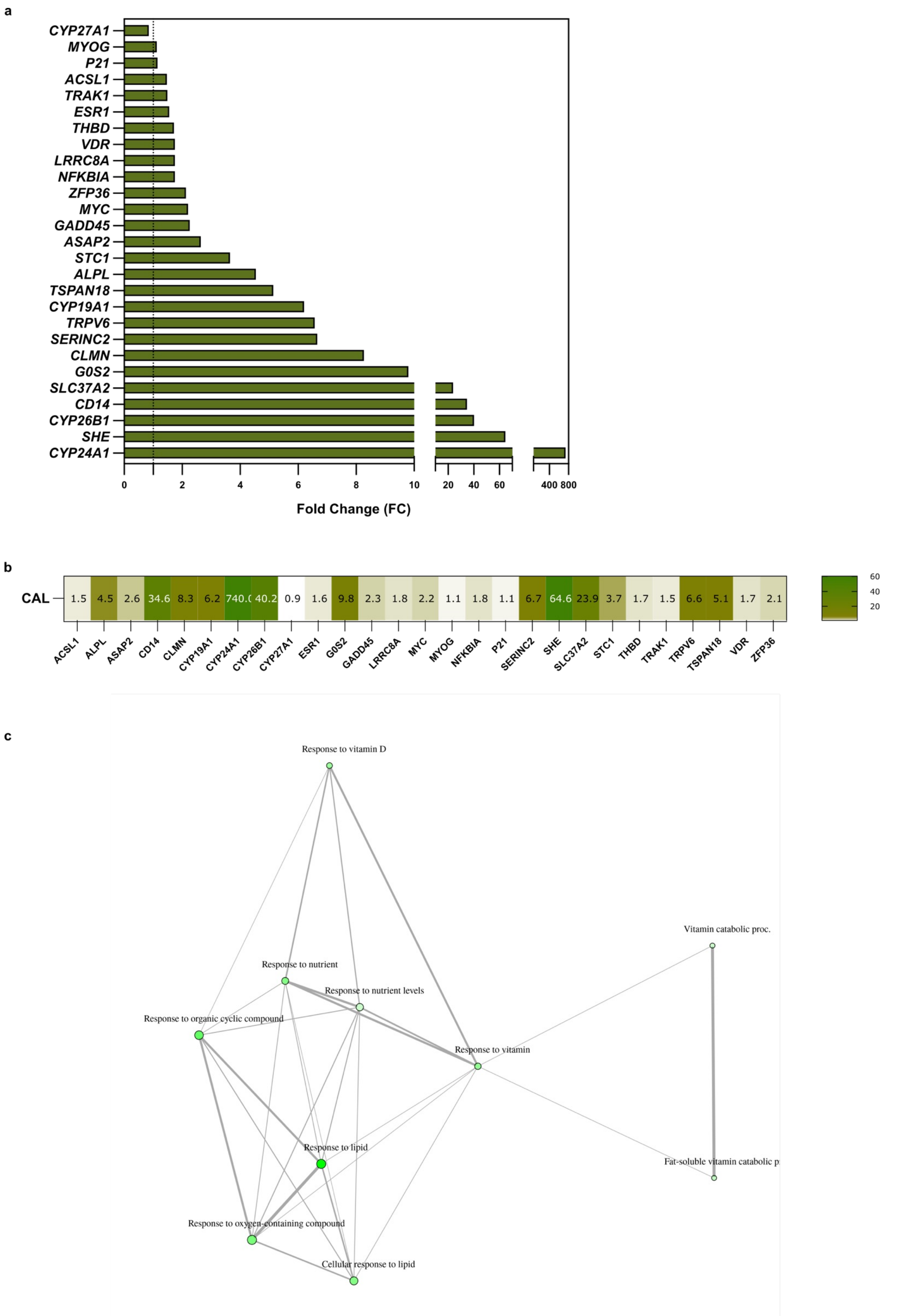


**Supplementary Fig. 12 | Calcitriol induces vitamin D receptor (VDR) target gene expression in DM1 3D muscle tissues.**
**a,b,** Fold-change (FC) values derived from RNA-seq analysis comparing untreated and calcitriol-treated DM1 3D muscle tissues. Genes shown correspond to previously reported primary vitamin D–responsive targets (Nurminen et al., 2019). **c,** Gene ontology (GO) network visualization generated using ShinyGO 0.85 based on differentially expressed genes following calcitriol treatment. Nodes represent enriched GO terms, with node size indicating the number of associated genes and edge thickness reflecting the proportion of shared genes between terms. Enrichment was performed using a cutoff of 0.4.

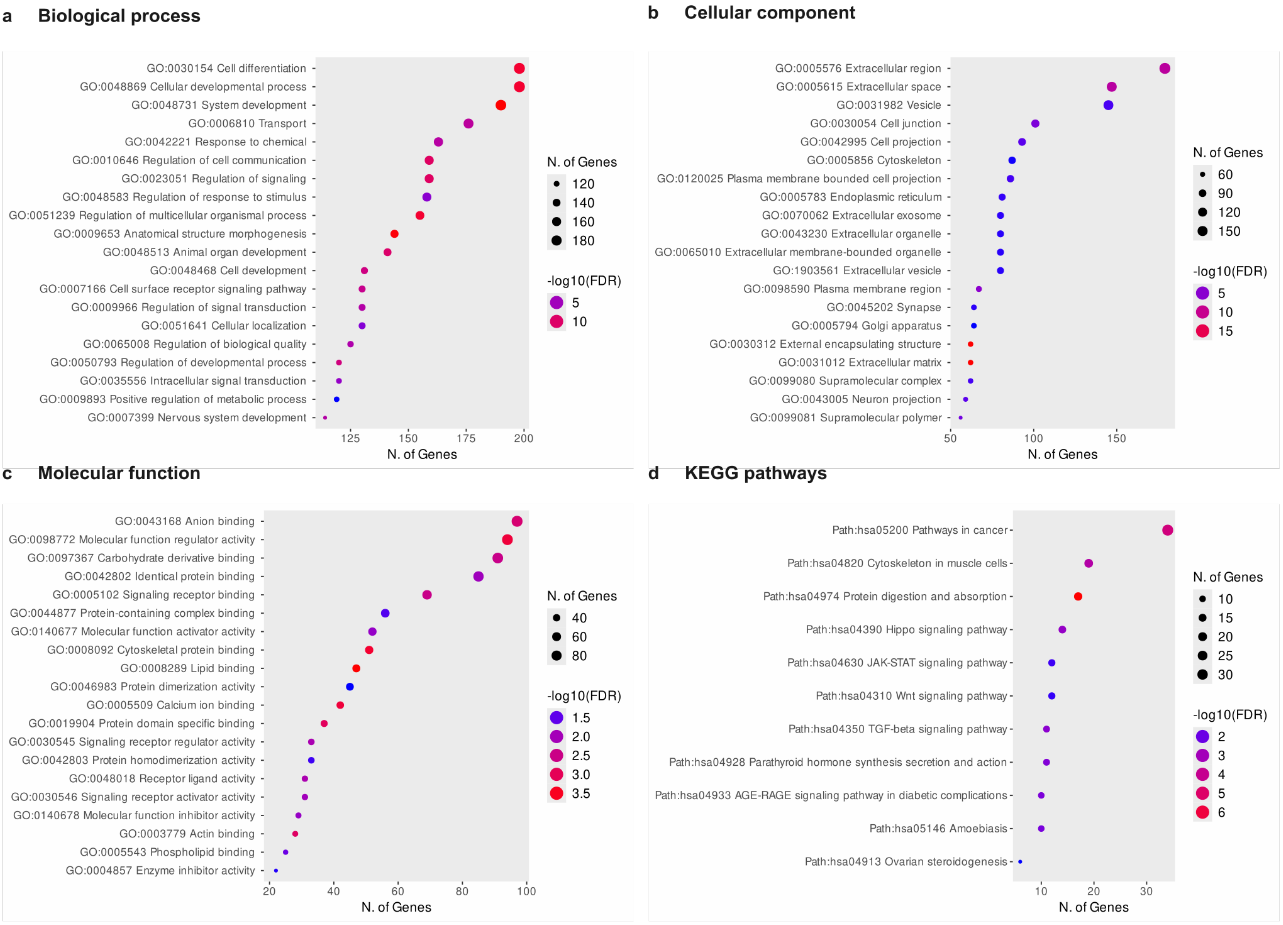


**Supplementary Fig. 13 | Functional enrichment analysis of differentially expressed genes in calcitriol-treated DM1 3D muscle tissues.** Gene Ontology (GO) and KEGG pathway enrichment analyses derived from RNA-seq data comparing calcitriol-treated (CAL) and untreated three-dimensional DM1 muscle tissues. Four annotation categories are shown: **(a)** Biological process, **(b)** Cellular component, **(c)** Molecular function, and **(d)** KEGG signaling pathways. Dot size indicates the number of genes associated with each term, and color intensity reflects statistical significance ($-\log_{10}$ FDR).

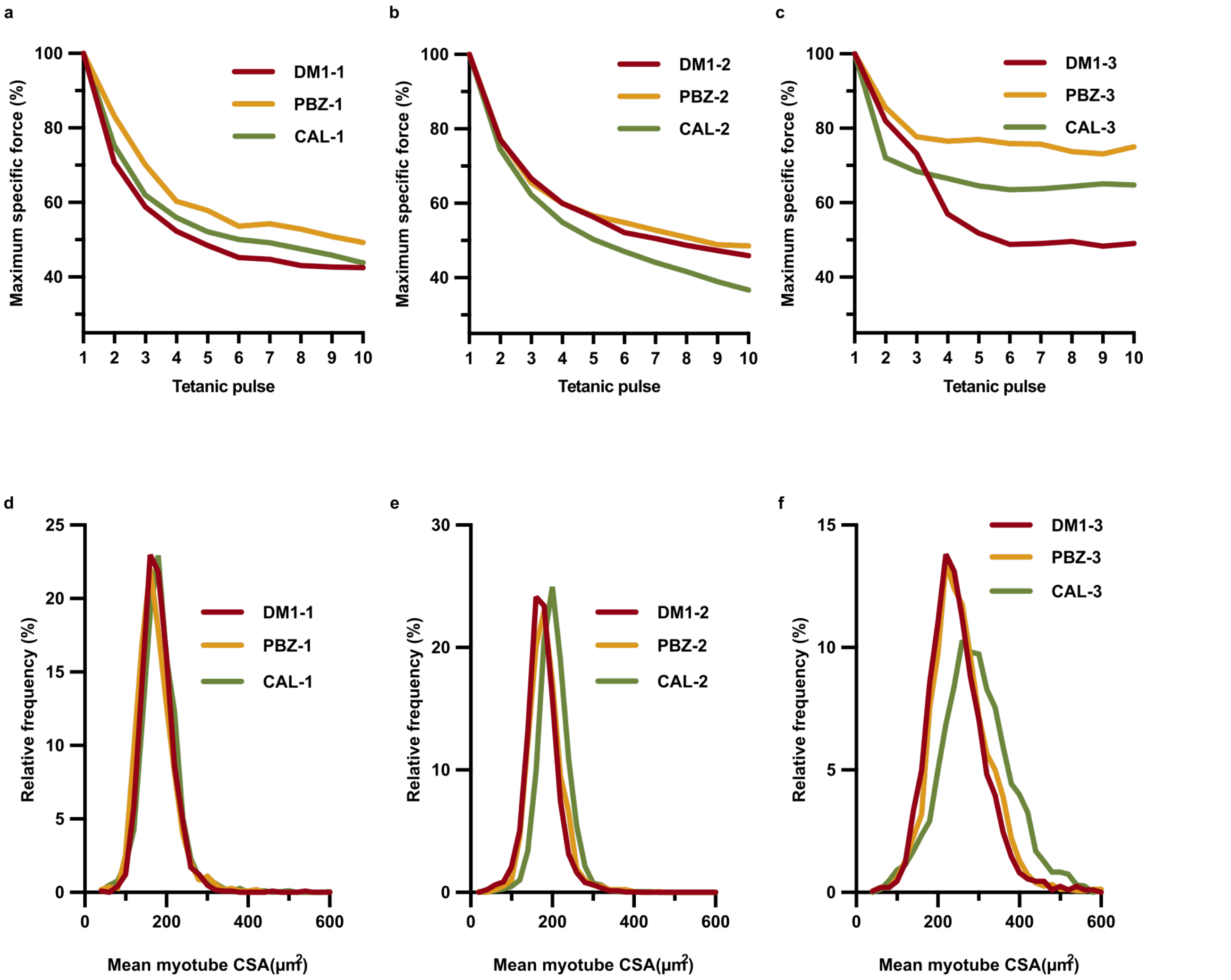


**Supplementary Fig. 14 | Effects of phenylbutazone and calcitriol on fatigue and myotube size in DM1 tissues.**
**a–c,** Normalized specific force (%) at each tetanic pulse relative to the first pulse, assessing fatigue behavior in DM1 tissues. No detectable changes were observed in **(a)** DM1-1 **(b)** DM1-2 tissues following treatment. In adult DM1-3 tissues **(c)**, PBZ- and CAL-treated samples exhibited altered force decay profiles compared with untreated DM1 tissues. **d–f,** Frequency distribution of myotube cross-sectional area (CSA, μm²) in DM1 tissues and following treatment with PBZ or CAL. CSA was measured from dystrophin-delimited membrane area in transverse cryosections (~800–900 myotubes per section). Frequency distributions are expressed as percentages of total myotubes. Data are shown as mean ± SEM. n = 5–8 tissues per condition.

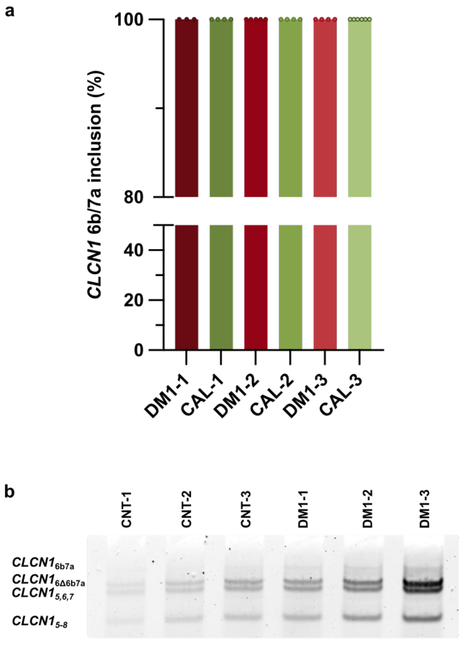


**Supplementary Fig. 15 | *CLCN1* splicing defects persist in DM1 3D muscle tissues following calcitriol treatment.**
**a,** Percentage of exon inclusion for *CLCN1* exons 6b/7a measured by semiquantitative RT–PCR in untreated DM1 and CAL-treated DM1 3D muscle tissues across the three patient lines. *GAPDH* was used as an internal control. **b,** Representative RT–PCR gels used for quantification in **(a)**.
